# Single-nucleus multi-omics reveals reversible molecular signatures in skeletal muscle of patients with heart failure with preserved ejection fraction

**DOI:** 10.64898/2026.10.07.756854

**Authors:** Maya Bose, Arushi Varshney, Nicholas J Carruthers, Peter Orchard, Mary Piper, Evanthia Pashos, Vladislav Dolgachev, Nandini Manickam, Erika Curka, Pierre M Jean-Beltran, Dalane W Kitzman, Eric B Fauman, Federico Damilano, Rachel J Roth Flach, Barbara Nicklas, Peter Brubaker, Stephen CJ Parker

## Abstract

Heart failure with preserved ejection fraction (HFpEF) is a complex systemic syndrome characterized by severe exercise intolerance. In addition to cardiac dysfunction, abnormalities in peripheral skeletal muscle are key contributors to reduced exercise capacity in HFpEF. Multiple studies show that exercise training and caloric restriction improve exercise capacity and quality of life in HFpEF patients. To map the HFpEF-associated molecular signatures in skeletal muscle and the effect of lifestyle intervention, we generated single-nucleus (sn) multiome (joint snRNA-seq and snATAC-seq) profiles of 140 biopsies. Our study included 95 individuals: 25 healthy controls and 70 HFpEF patients, of whom 45 had paired biopsies collected before and after a supervised diet and exercise intervention. From 132,568 nuclei, we resolved 15 skeletal muscle cell types and identified HFpEF-specific remodeling, including type 1 fiber and pericyte depletion and type 2x fiber enrichment, reflecting the cellular basis of exercise intolerance. Transcriptional changes in HFpEF muscle indicated dysregulation of metabolic pathways, stress-responsive programs, cytoskeletal integrity, and calcium signaling. Transcription factor (TF) motifs enriched in differentially accessible chromatin corresponded to concordantly differentially expressed TF-genes, revealing coordinated regulatory programs underlying these changes. Importantly, HFpEF-related changes in both chromatin accessibility and transcription were partially reversible following diet and exercise. Integrative multiomic velocity analyses revealed increased coordinated repression in both chromatin accessibility and transcription of calcium handling genes in HFpEF. HFpEF-linked chromatin was enriched for HF and cardiometabolic GWAS signals, and cross-modal analyses further nominated candidate causal variants and effector genes linking genetic risk to peripheral muscle dysfunction in HFpEF. Together, these findings highlight the molecular mechanisms underlying skeletal muscle remodeling in HFpEF and identify pathways responsive to lifestyle interventions.

## Introduction

Heart failure (HF) is a major cardiovascular condition associated with high morbidity and mortality, which is predicted to affect upwards of 6 million people in the US^1^. Heart failure with preserved ejection fraction (HFpEF) is a clinical subtype of HF characterized by structural and functional abnormalities of the heart but with a preserved left ventricular ejection fraction (≥50%)^2,3^. About half of all HF cases are HFpEF, the other half being heart failure with reduced ejection fraction (HFrEF). HFpEF is particularly common in older persons and females, in whom it accounts for >80% of HF cases^4–10^. The prevalence of HFpEF is increasing disproportionately as a result of the aging populace and increased occurrence of diabetes and obesity^11^, and has recently become recognized as a systemic, multi-organ disorder with frequent metabolic comorbidities, particularly overweight / obesity which is present in >80% of patients with HFpEF.

HFpEF is associated with poor quality of life, increasing healthcare costs, significant morbidity, and premature mortality^12^. Patients with HFpEF have severe exercise intolerance with symptoms of exertional dyspnea and fatigue. Exercise intolerance is measured objectively as severely reduced peak oxygen uptake during peak exercise (peak VO_2_)^13–16^, and contributes to functional limitation and impaired daily living. Our work and others’ support that exercise intolerance in HFpEF arises not only from impaired cardiac output but also from abnormalities in peripheral tissues, particularly skeletal muscle^10,17^. These abnormalities include muscle atrophy, accumulation of intermuscular fat^18^, reduced oxidative capacity, and altered metabolism, along with impaired skeletal muscle vasculature marked by reduced capillarization^19^. Several studies have demonstrated that HFpEF patients have significant depletion in type 1 muscle fibers and an expansion in glycolytic type 2 fibers when compared to healthy, age-matched controls^15^.

Healthy type 1 fibers display high mitochondrial density and contribute to aerobic endurance, which appears impaired in HFpEF. Depletions in muscle mass have been observed in a mouse model of heart failure with reduced ejection fraction (HFrEF) even when physical conditioning is maintained^14,20,21^. Likewise, exercise intolerance due to muscle dysfunction is observed in mouse and rat models of HFpEF^22–24^. Collectively, these observations implicate skeletal muscle dysfunction as a key contributor to exercise intolerance in HFpEF, underscoring the need to better understand its molecular mechanisms.

Despite its importance and increasing prevalence, treatment options for HFpEF remain limited. Exercise training and caloric restriction diet were the first two treatments shown to improve outcomes, including exercise capacity (peak VO_2_) and quality of life in patients with HFpEF^10^, and appear to be mediated primarily by extracardiac mechanisms, such as improvements in body composition and skeletal muscle function^25,26^. Similar improvements in skeletal muscle due to exercise training have been observed in a HFpEF mouse model^27^. These findings suggest that interventions improving HFpEF outcomes may act, at least in part, through peripheral adaptations in skeletal muscle. However, the molecular mechanisms driving these improvements in humans remain unclear.

To better understand the skeletal muscle dysfunction underlying reduced exercise capacity in HFpEF and the molecular effects of diet and exercise interventions, we obtained 140 skeletal muscle biopsies from 95 human participants, including HFpEF patients, healthy, age-matched controls, and HFpEF patients undergoing supervised diet or exercise/resistance training interventions. For 45 participants with HFpEF, matched biopsy samples were collected before and after intervention, enabling direct characterization of molecular changes associated with improved exercise capacity. We performed paired single-nucleus (sn) multiome profiling (snRNA-seq and snATAC-seq) on these biopsies using a randomized block study design in which nuclei from multiple samples were pooled into batches balanced for age, sex, cohort, and disease status, followed by genotype-based demultiplexing. This integrated multiomics approach enabled simultaneous characterization of cell-type composition, transcriptional programs, and regulatory chromatin landscapes underlying HFpEF skeletal muscle dysfunction and their exercise-associated partial restoration toward a non-disease state.

## Results

### Patient samples

We acquired frozen *vastus lateralis* skeletal muscle tissue biopsies from Wake Forest University School of Medicine. Biopsies were originally obtained as part of the SECRET, and SECRET II clinical trials, in addition to a healthy control (HC) cohort of age-matched subjects^25,28^. In the SECRET I cohort, participants diagnosed with HFpEF provided biopsies after a 20-week supervised diet (n=5), exercise (n=7), combined diet and exercise (n=10), or no intervention (n=3). The SECRET II cohort (n=45) included participants diagnosed with HFpEF, from whom paired biopsies were collected both before and after a supervised diet and exercise intervention. The participants in the healthy, age-matched control cohort were free of heart failure and other chronic medical conditions. In total, we obtained 140 muscle biopsies from 95 participants (71% female) (Fig. 1a). The age, sex, and BMI of participants are reported in Supplemental Table S1.

**Figure 1:**
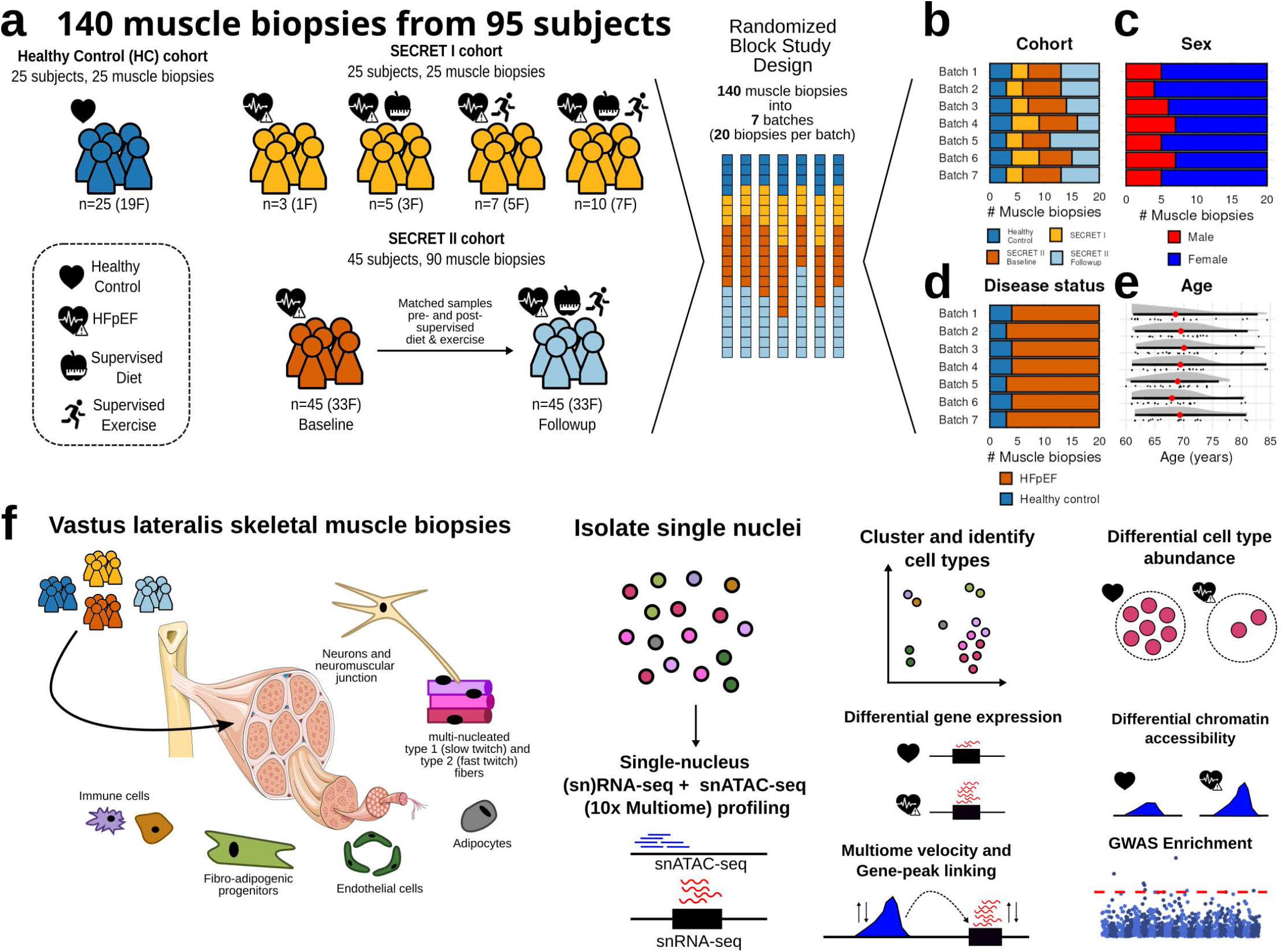
Single-nucleus multi-omic profiling of 140 skeletal muscle biopsies. (a) Overview of cohort participants. Biopsies were obtained from 3 cohorts: SECRET 1, SECRET II, and a cohort of age-matched healthy controls. The healthy control (HC) cohort contributed one biopsy from each of 25 healthy participants who received no treatment. The SECRET I cohort contributed one biopsy each from 25 participants diagnosed with HFpEF, assigned to one of four treatment conditions: no treatment, supervised exercise, supervised diet, or combined supervised diet and exercise. The SECRET II cohort contributed two muscle biopsies each from 45 participants diagnosed with HFpEF, collected before and after a supervised diet and exercise intervention. All samples were multiplexed into 7 batches of 20 samples each, designed to ensure an approximately even distribution of (b) cohort, (c) biological sex, (d) disease status, and (e) age across batches (bar with red point indicates mean quantile interval). (f) Schematic of the experimental and analytical workflow. Skeletal muscle biopsies were collected from the vastus lateralis of each participant. Single nuclei were isolated from all samples and profiled using paired snRNA-seq and snATAC-seq on the 10x Genomics Chromium platform. Downstream analysis included clustering and cell type identification, differential cell type abundance, differential gene expression, differential chromatin accessibility, gene-peak linking, and functional GWAS enrichment.

### Single-nucleus multiome profiling of skeletal muscle with randomized block batch assignment

To profile gene expression and chromatin accessibility, we performed paired snRNA-seq and snATACseq on nuclei isolated from the skeletal muscle biopsies using the 10x Genomics Chromium platform (Fig. 1f). We processed the samples in 7 batches of 20 biopsies each, designing the batches to evenly distribute cohorts, sex, HFpEF status, and participant age (Fig. 1b-e) across batches. This design allowed us to attribute variation between batches to technical rather than biological differences.

We obtained 102,941 high-quality multiome nuclei after quality control (QC) and filtering on both ATAC and RNA modalities (Fig 2a). For RNA-based clustering we included an additional 29,627 nuclei that passed all RNA QC thresholds, and 61,565 nuclei from additional 5’ snRNA sequencing of the same samples. Downstream analyses were performed using nuclei from the multiome modality. We performed integration across batches using the Harmony approach implemented in Seurat^29^. This analysis identified 15 clusters, which we annotated using known marker gene expression (Fig. 2c, Extended Data Fig 1, Supplemental Dataset 1). These marker genes broadly recapitulated cluster specific patterns in the corresponding chromatin accessibility profiles (Fig. 2d).

**Figure 2:**
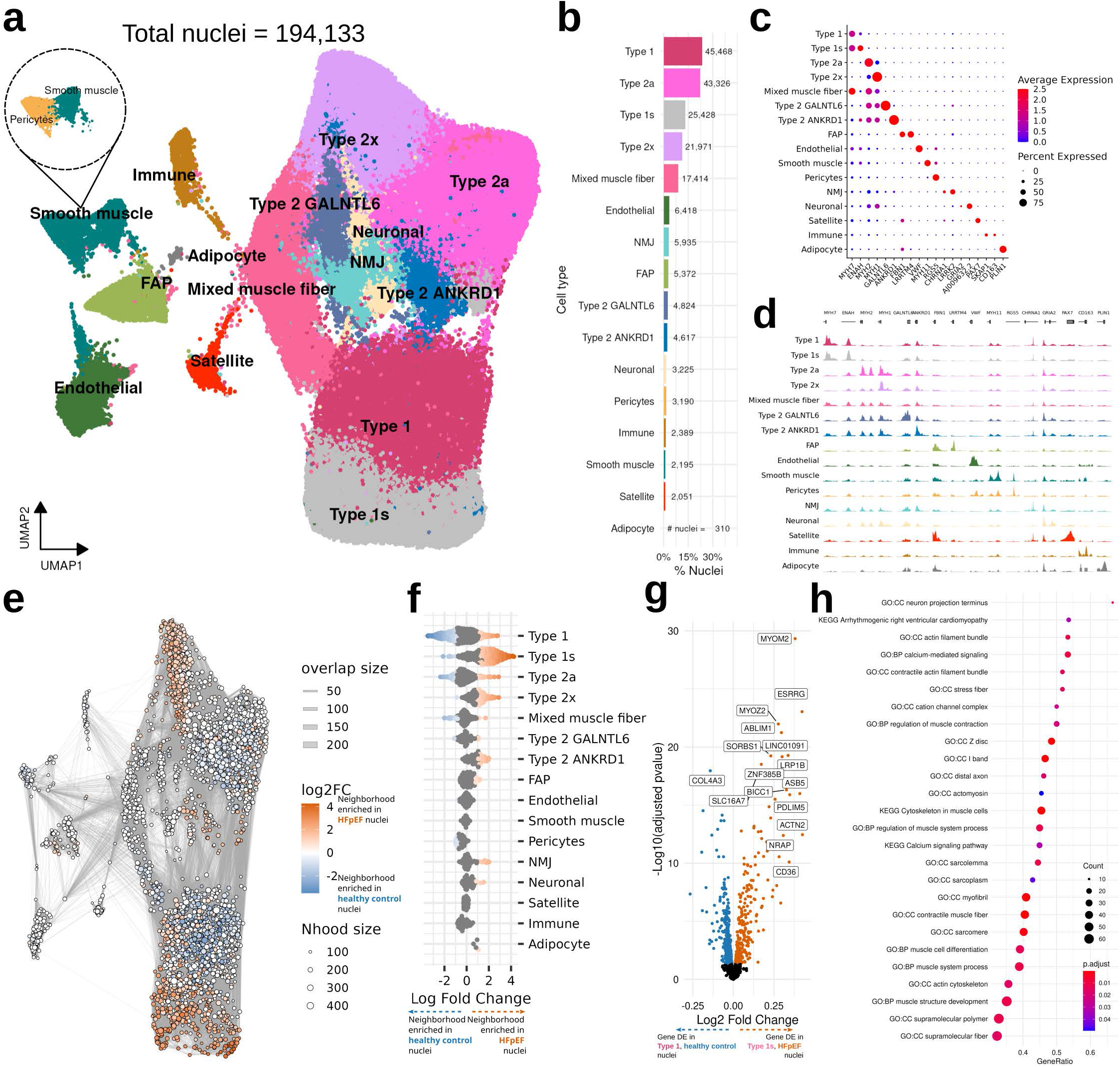
Cell type clustering and differential abundance analysis reveal a muscle fiber transition linked to calcium-mediated signalling. (a) UMAP (Uniform Manifold Approximation and Projection of 194,133 skeletal muscle nuclei profiled by RNA expression, derived from multiome and 5’ RNA sequencing. Sub-clustering of the smooth muscle cluster was performed to resolve a distinct pericyte population. (b) Nuclei counts and relative proportions of each identified cell type cluster. (c) Marker gene expression and (d) chromatin accessibility at marker gene promoters across cell type clusters. (e) Neighborhood graph of nuclei colored by differential abundance between healthy controls and HFpEF participants. Orange neighborhoods are significantly (Spatial FDR < 0.05) enriched in HFpEF nuclei; blue neighborhoods are significantly enriched in healthy control nuclei. Line thickness between neighborhoods reflects the degree of overlap in nuclei. (f) Beeswarm plot grouping neighborhoods by their majority cell type. Orange neighborhoods are significantly (Spatial FDR < 0.05) enriched in HFpEF nuclei, while blue neighborhoods are significantly enriched in healthy nuclei. (g) Volcano plot showing differential gene expression results between Type 1 nuclei in healthy-enriched neighborhoods (blue) and Type 1s nuclei in HFpEF-enriched neighborhoods (orange). Colored points are significant at FDR < 5%. (h) Gene Ontology (GO) terms significantly enriched (FDR < 5%) among genes differentially expressed (FDR < 1%) between Type 1s nuclei in HFpEF-enriched neighborhoods and Type 1 nuclei in HFpEF-depleted neighborhoods.

Consistent with the multinucleated nature of skeletal muscle fibers, nuclei in our data were predominantly derived from myofibers, including type 1, type 2a, and type 2x, and mixed muscle fiber subtypes. We observed two *MYH7* expressing clusters, annotated as type 1 and type 1s, with the latter clustering separately from the primary type 1 fiber population. We also detected two additional type 2 fiber subtypes that showed *MYH1* and *MYH2* expression along with higher levels of *GALNTL6* and *ANKRD1.* We identified endothelial, neuro-muscular junction (NMJ), smooth muscle, fibro-adipogenic progenitors (FAPs), neuronal, immune, satellite, and adipocyte cell types that varied in abundance (Fig. 2b). Further sub-clustering identified fine-grained cell types, separating pericyte cell type from the smooth muscle cluster (Fig 2a), macrophages and T cells from immune cells, and identified distinct functional subtypes of FAP and endothelial cells (Extended Data Fig 2). These results highlight the utility of our dataset as a resource for resolving skeletal muscle cell type heterogeneity.

### Type 1s and Type 1 clusters demonstrate disease-correlated differential abundance with differences in calcium-mediated signaling

HFpEF is associated with decreased vasculature and slow-twitch type 1 muscle fibers, and increased fast-twitch type 2x fibers^15^. To test for these compositional changes and investigate their molecular basis in our data, we performed differential abundance analysis on the multiome nuclei using Milo^30^. Milo constructs a k-nearest neighbors graph of nuclei, groups them into cluster-agnostic neighborhoods based on gene expression similarity, and tests these neighborhoods for differential abundance between conditions using a negative binomial generalized linear model (Fig. 2e). Neighborhoods are then assigned to their majority-represented cluster to enable comparisons between cell types.

Differential abundance analysis revealed a statistically significant (spatial FDR < 5%) increase in HFpEF nuclei in Type 2x neighborhoods, and a statistically significant decrease in HFpEF nuclei in pericytes, which recapitulates previously reported compositional shifts in HFpEF skeletal muscle. Notably, neighborhoods representing the Type 1s subcluster were significantly enriched in HFpEF, while neighborhoods representing the Type 1 cluster were significantly depleted in HFpEF (Fig. 2f). This suggested that the type 1s subcluster represents a remodeled form of type 1 muscle fiber that is more abundant in HFpEF patients. Cell type abundance changes in HFpEF patients following supervised diet and exercise (HFpEF-followup) as compared to baseline (HFpEF) were more modest, with some significant depletion of Type 1s and Type 2a fibers following intervention (Extended Data Fig 3).

To assess transcriptional differences between type 1 and type 1s clusters, we focused on type 1 neighborhoods enriched in healthy controls and type 1s neighborhoods enriched in HFpEF, and performed a differential expression (DE) analysis (marker gene identification) in Milo (Fig. 2g, Supplemental Table S2). Pathway analysis of these marker genes revealed an enrichment in calcium-mediated signaling and regulation of muscle system processes (Fig. 2h, Supplemental Table S3). Calcium-mediated signaling plays a central role in muscle fiber contraction and differentiation, and perturbations in this pathway can strongly affect muscle fiber function^31,32^.

These findings suggest that the type 1s fibers enriched in HFpEF, while still identifiable as slow twitch based on MYH7 expression (Fig. 2c), are metabolically dysregulated in a manner that may impair energy efficiency, thus contributing to weakness and exercise intolerance. Notably, HFpEF type 1s muscle fibers showed increased expression of *AKAP6*^33^, *CAMK2D*^34–38^, *CASQ2*^39–41^, *CCR3*^42^, *ITPR2*^43,44^, *MCU*^45^, and *MYH7B*^46,47^ consistent with gene signatures of cardiac hypertrophy. This suggests that calcium-mediated signaling dysregulation, which drives hypertrophy in the heart, may similarly compromise skeletal muscle function. HFpEF type 1s muscle fibers also showed decreased expression of NFAT-calcineurin signaling genes *NFATC3* and *PPP3CA*, the reverse of what is observed in cardiac hypertrophy^48^. The observed concurrent increase in *CAMK2D* expression and decrease in *NFATC3* and calcineurin (*PPP3CA/B*) expression indicates a transition from slow twitch, type 1 fibers towards a less energy-efficient, more type 2-like phenotype^49,50^.

### Differentially expressed genes in HFpEF skeletal muscle are enriched in heart failure pathway genes, revealing divergent symptoms of shared transcriptional dysregulation

To understand the molecular mechanisms underlying the improved quality of life and peak VO_2_ observed in HFpEF-followup patients, we performed differential gene expression analysis across all profiled cell types. We modeled gene expression differences between healthy controls and HFpEF, and between HFpEF and HFpEF-followup, using MAST^51^ with age, batch, sex as fixed-effect variables and sample ID as a random effect variable to control for within-sample correlation. Compared to pseudobulk approaches such as DESeq2^52^, MAST allowed us to better leverage the resolution of our single-nucleus data (Extended Data Fig 4). To further increase statistical power and interpretability, we merged the Type 1 and Type 1s clusters into a single Type 1 cluster (Fig. 3a).

**Figure 3:**
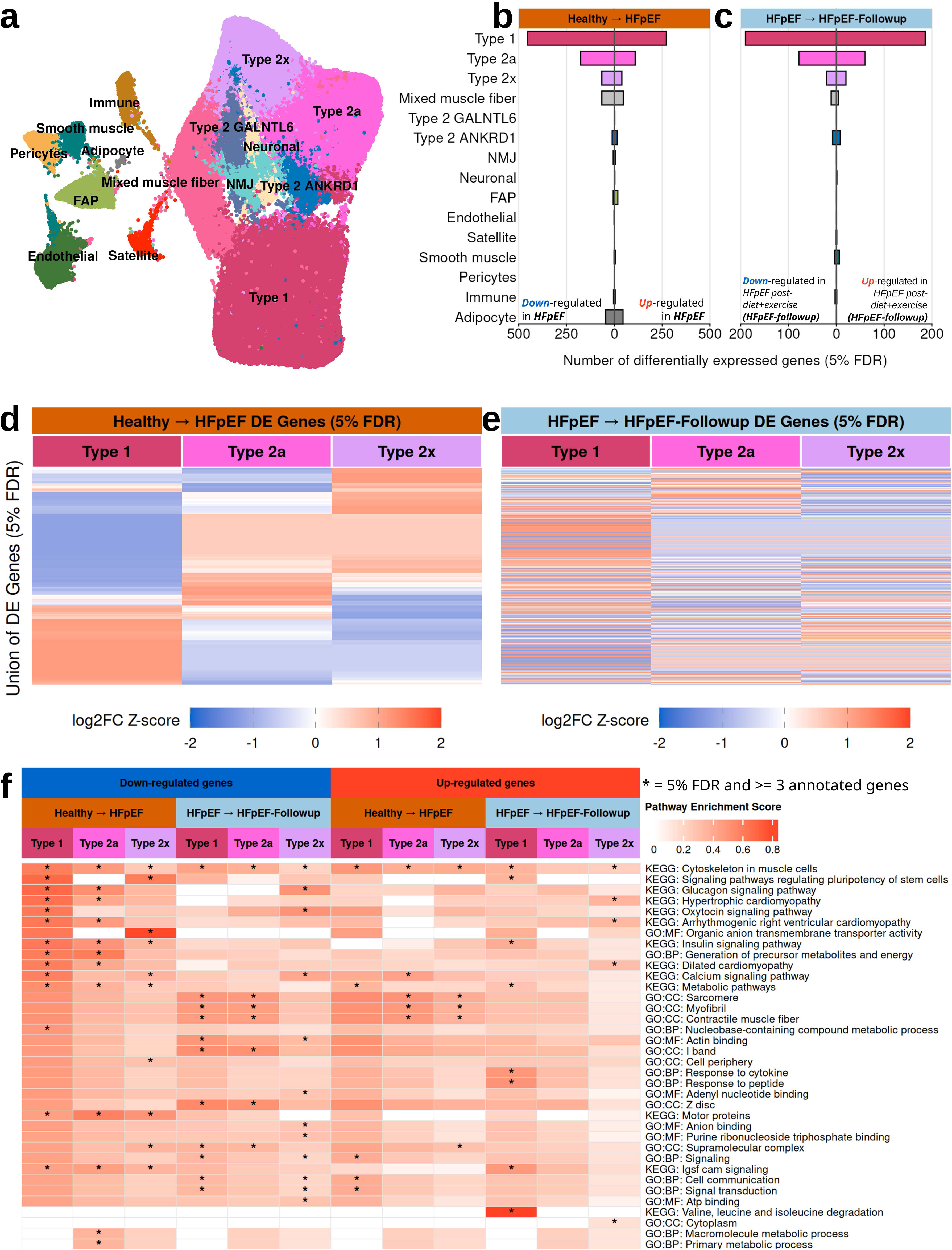
Differential gene expression analysis across cell types share transcriptional responses to HFpEF and supervised diet and exercise intervention across major muscle fiber types. (a) UMAP of skeletal muscle nuclei with Type 1 and Type 1s clusters merged to increase statistical power for differential expression analysis. Number of significant differentially expressed genes (FDR < 5%) per cell type, shown separately for the (b) healthy-to-HFpEF and c) HFpEF-to-HFpEF followup (post-supervised diet and exercise) comparisons. Bars extending to the right indicate genes upregulated in HFpEF or HFpEF-followup; bars extending to the left indicate genes downregulated in those states. Heatmaps of log2 fold change (Z-Score scaled) for the union of significantly differentially expressed genes across the three major muscle fibers (Type 1, Type 2x, and Type 2a) for (d) the healthy-to-HFpEF and (e) HFpEF-to-HFpEF-followup comparisons. Each row represents one gene; rows are clustered by log2FC in the healthy-to-HFpEF comparison in both heatmaps. (f) Pathway enrichment heatmap showing the top 10 most significant pathways for up-regulated and down-regulated genes across each cell type and comparison. Color intensity reflects the pathway enrichment score (PES), defined as the ratio of input genes that are annotated to a given pathway term to all genes annotated to that term. Asterisks (*) indicate pathways significant at FDR < 5% with at least 3 annotated genes.

Across all cell types, we identified 1,698 significantly (FDR < 5%) DE genes between healthy controls and HFpEF (Fig. 3b, Supplemental Table S4), and 807 DE genes between HFpEF and HFpEF post-diet and exercise (HFpEF-followup) (Fig. 3c, Supplemental Table S5) across all cell types. We noted that the majority (60%) of DE genes in the healthy-to-HFpEF comparison are down-regulated, suggesting that the observed impairment in muscle endurance observed in HFpEF patients is driven primarily by loss-of-function.

To characterize the pathways enriched among DE genes in the three major muscle fiber types, we performed gene set enrichment analysis with Gene Ontology (GO) and KEGG terms (Fig. 3f, Supplemental Tables S6, S7). Genes upregulated in HFpEF relative to healthy controls were consistently enriched for cytoskeleton in muscle cells across all three fiber types, and for structural muscle constituents (e.g. sarcomere, myofibril, and contractile muscle fiber) in Type 2a and Type 2x fibers. Simultaneously, we see dysregulation of oxytocin and calcium signaling pathways in genes downregulated in HFpEF Type 1 fibers relative to healthy controls. As previously discussed, these pathways are key drivers of muscle fiber differentiation and development^32,53,54^. Genes down-regulated in HFpEF were enriched for terms associated with heart disease, including arrhythmogenic right ventricular cardiomyopathy, hypertrophic cardiomyopathy, and dilated cardiomyopathy, in Type 1 and Type 2a fibers. This pattern suggests that the same transcriptional dysregulation that drives cardiac hypertrophy in HFpEF may simultaneously promote skeletal muscle atrophy.

### DE Genes related to muscle development in HFpEF reverse direction after supervised diet and exercise

Comparing the direction of effect between the healthy-to-HFpEF and HFpEF-to-HFpEF followup contrasts revealed a striking pattern of transcriptional reversal. We observed that genes upregulated in HFpEF relative to healthy controls tended to decrease in expression following the intervention, while genes downregulated in HFpEF tended to recover toward healthy levels (Fig. 3d,e). This trend was apparent both at the level of individual genes and at the pathway level, where gene sets significantly enriched among upregulated genes in one comparison were correspondingly enriched among downregulated genes in the other, and vice versa (Fig. 3f).

Among genes with FDR < 5% in one comparison and at least nominal significance (p < 0.05) in the other, the vast majority showed opposing directions of effect between the two contrasts (∼87% overall, 87% in Type 1, 86% in Type 2a, 89% Type 2x) (Fig. 4a, Extended Data Fig 5a). We show the top 10 reversing genes in the three fiber types in Fig 4b. Absolute effect sizes from HFpEF to HFpEF-Followup were relatively smaller than the effects of disease (Extended Data Fig 5b). Taken together, these findings show that, at the level of gene transcription, supervised diet and exercise are able to partially reverse the molecular signatures of HFpEF in skeletal muscle.

**Figure 4:**
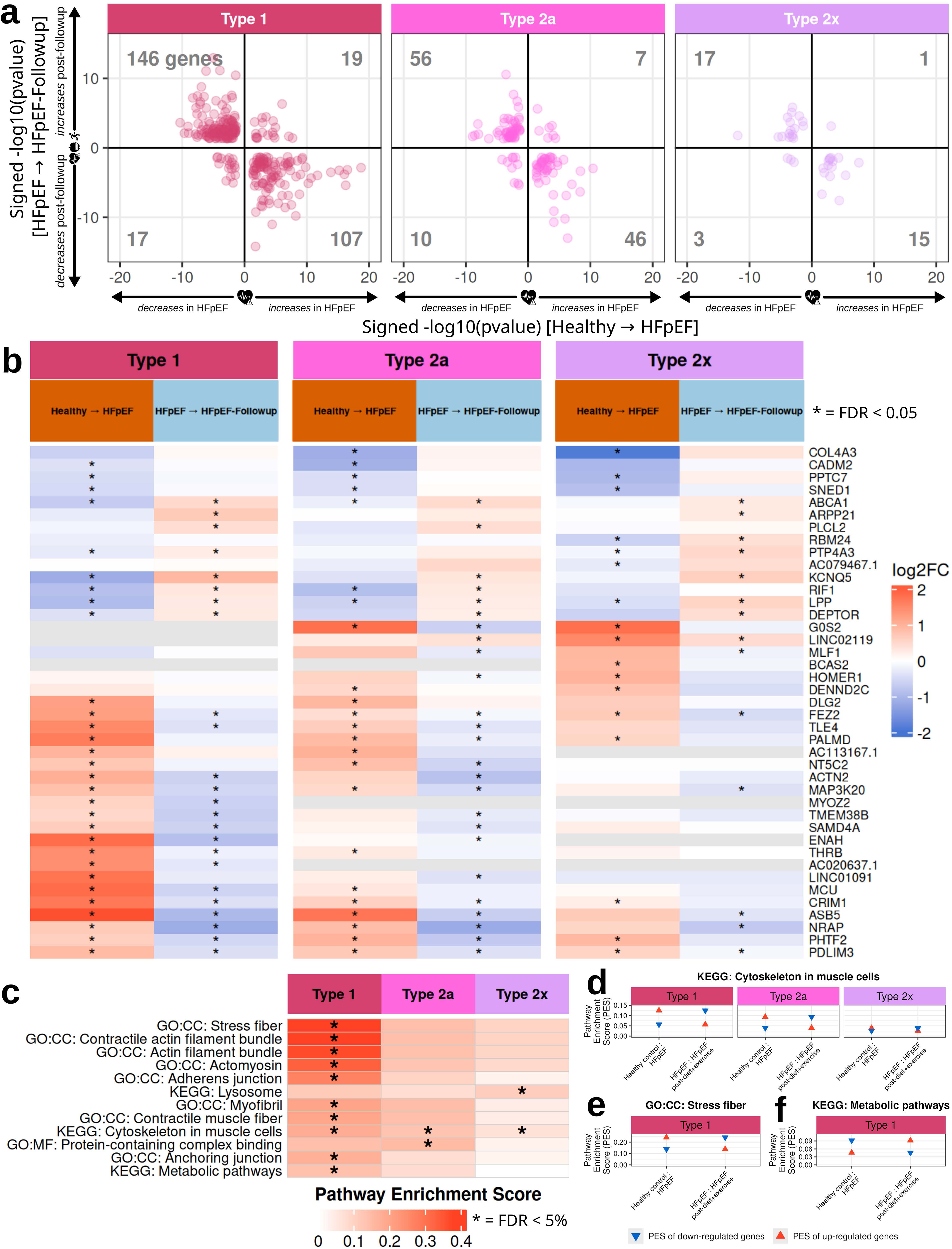
Supervised diet and exercise intervention partially reverses the transcriptional signature of HFpEF across muscle fiber types. (a) Scatter plots comparing signed-log10(p-value) of significantly differentially expressed between the healthy-to-HFpEF (x-axis) and HFpEF-to-HFpEF-followup (y-axis) comparisons for Type 1, Type 2a, and Type 2x muscle fibers. Genes are included if they reach FDR < 5% in one comparison and at least nominal significance (p < 0.05) in the other. Genes in the upper-left and lower-right quadrants show expression changes that are reversed by the supervised diet and exercise intervention. Numbers in each quadrant indicate the count of significant genes. (b) Heatmap of log2FC for the 10 reversing genes with the largest absolute log2 fold change each cell type (for the three major muscle fibers) and contrast. Asterisks (*) indicate significance at FDR < 5%. (c) Pathway enrichment heatmap of the top 10 significantly enriched Gene Ontology (GO) and KEGG pathways (FDR < 5%) among reversal genes across the three muscle fiber types. Color intensity reflects the pathway enrichment score (PES), defined as the ratio of input genes that are annotated to a given pathway term to all genes annotated to that term. PES for down-regulated (blue triangle) and up-regulated (red triangle) reversal genes across the healthy-to-HFpEF and HFpEF-to-HFpEF-followup comparisons are shown for (d) Stress fiber GO Cellular Component (CC) term, (e) cytoskeleton in muscle cells KEGG term, and (f) metabolic pathways KEGG term.

Reversal genes across all three major muscle fibers were enriched for the cytoskeleton in muscle cells KEGG pathway, with a majority of genes transitioning from up-regulation in HFpEF to down-regulation following intervention (Fig. 4c, d, Supplemental Table S8). This included significantly increased expression of *NRAP* and *PDLIM3* across the three muscle fiber types, and *ACTN2, ANKRD2, CSRP3,* and *MYOZ2* specifically in Type 1 fibers. Collectively, these genes stabilize sarcomere structure and transmit mechanical tension while under remodelling stress^55,56,56–66^. Conversely, genes involved in filament maintenance and calcium-ion coupling, including *TMOD1, COL4A3, ACTN2,* and *OBSCN*, were suppressed in HFpEF and restored after the intervention^65,67–75^. Variants in several of these genes have previously been associated with cardiac disease, suggesting that shared transcriptional dysregulation may produce distinct symptoms in skeletal and cardiac muscle^59,76^.

A related pattern emerged in the stress-fiber GO cellular component gene set in Type 1 fibers (Fig. 4c,e). In HFpEF, genes associated with actin networks and stress fiber formation, including *ABLIM1*^77^, *SORBS1*^78,79^ and *PDLIM3/5*^80^, were upregulated, while genes that normally promote muscle hypertrophy, including PXN and *LPP*, were down-regulated^81–83^. *SORBS1* is a key adaptor protein in the insulin signalling pathway, connecting the cytoskeletal remodeling observed in HFpEF skeletal muscle to the broader metabolic disruption characteristic of insulin and type 2 diabetes^84,85^.

Following this observation, reversal genes in Type 1 fibers were also enriched for metabolic pathway KEGG term, with a majority of genes being suppressed in HFpEF followed by recovery in HFpEF-followup (Fig. 4c,f). Relevant genes included those contributing to mitochondrial-oxidative capacity (*CS*^86–88^, *NNT*^89,90^, *MDH1*^91,92^*, G0S2*^93^*, MCU*^94,95^), branched-chain amino acid catabolism (*BCKDHB*^96,97^, *MCCC1*^98^, *MCEE*^99,100^*, DEPTOR*^101^), glucose uptake (*ABCA1*^102^), and CoA biosynthesis (*PANK1*^103^), all of which are associated with insulin resistance^104–106^. This transcriptional profile parallels the metabolic disruption observed in T2D, and its partial reversal following the supervised diet and exercise intervention provides encouraging evidence that therapeutic strategies may be able to restore skeletal muscle metabolic function in HFpEF patients.

### snATAC-seq reveals disease and followup-associated chromatin remodeling

We leveraged our snATAC data to assess the effects of HFpEF, and supervised diet and exercise follow-up on chromatin accessibility in skeletal muscle cell-types. To maximize power in the sparse snATAC modality, we considered ‘coarse’ cell-type cluster annotations, and merged type 1 and type 1s into type 1, and type 2a, 2x, GALNTL6, and ANKRD1 into a single type 2 fiber cluster (Fig 5a). We called reproducible peaks in these 11 clusters across batches and harmonized coordinates across clusters to obtain 203,529 peaks (Methods). PCA on pseudobulk sample counts for type 1 fiber peaks reflected differences in chromatin accessibility across samples by disease condition (Fig 5a).

**Figure 5:**
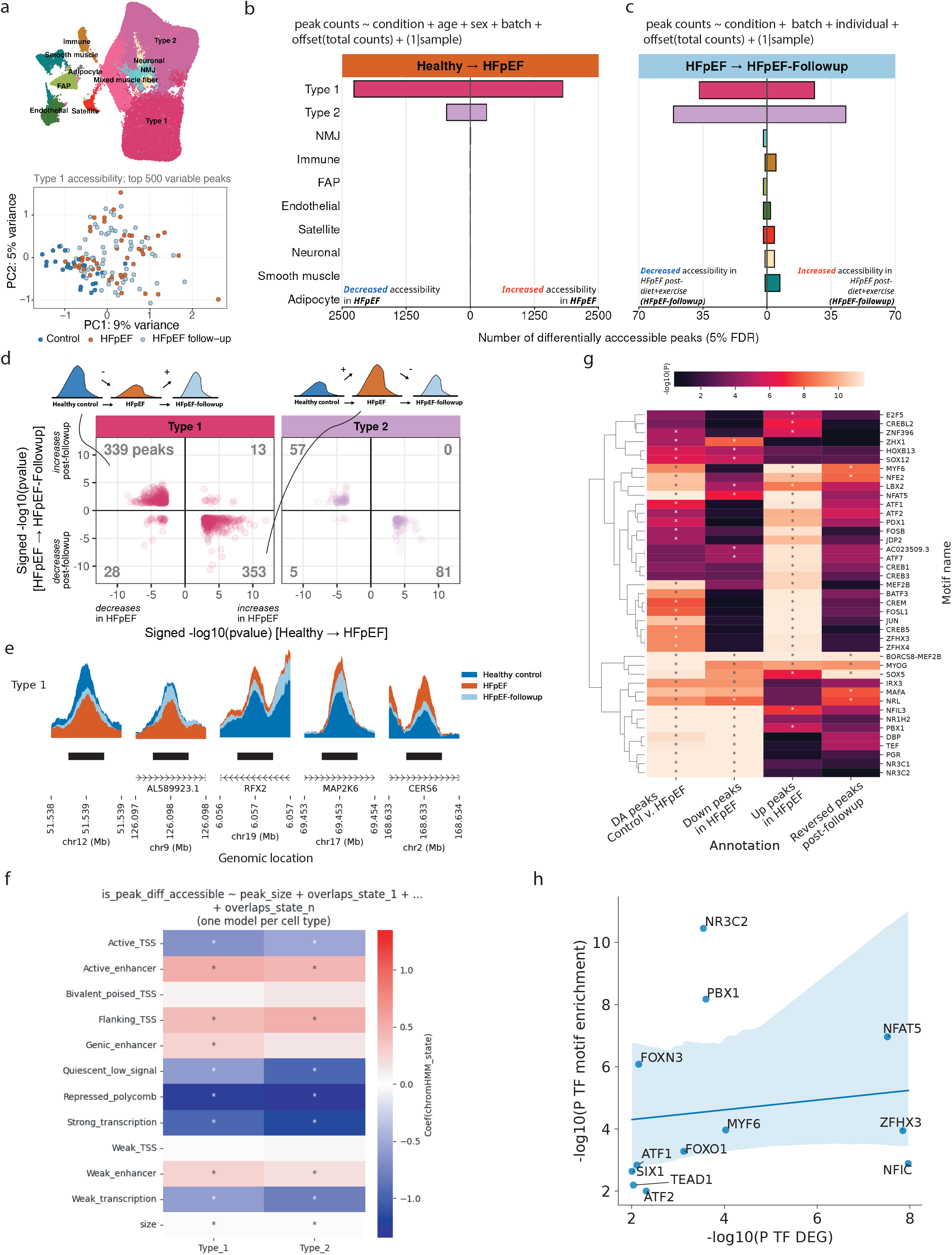
snATAC-seq reveals disease and followup -associated chromatin remodeling. a) ‘Coarse’ cluster annotations are considered to optimize power in snATAC analyses. PCA plot for type 1 fiber peak pseudobulk sample counts. Number of DA peaks identified (FDR < 5%) across clusters in healthy control v. HFpEF (b) and HFpEF v. HFpEF-follow-up (c). d) DA peak direction of effect comparison between the two contrasts for type 1 and type 2 fibers. e) ATAC-seq signal tracks by cluster and condition showing a cell-type-specific DA peak at the *MAP2K6* gene locus. f) Chromatin state overlap enrichment for muscle fiber DA peaks. (*) shows significant enrichment after Bonferroni correction. g) Motif enrichment in type 1 fiber DA peaks, peaks increasing and decreasing in accessibility in HFpEF and peaks reversing after followup. h) TF gene DE significance vs the enrichment of the cognate DNA-binding motif in DA peaks.

We next tested for differential accessibility (DA) in clusters by disease condition. Exploring pseudobulk and nucleus-level approaches (Extended Data Fig 6 a-c), we optimized DA testing in a nucleus-level negative binomial generalized linear mixed model (NB-GLMM), with sample included as a random effect to account for the non-independence of nuclei originating from the same individual. Comparing healthy controls with HFpEF nuclei, we identified the most DA peaks (FDR < 5%) in type 1 (4,072) and type 2 (782), with more peaks showing decreased accessibility in HFpEF (Fig 5b, Supplemental Dataset 2). We detected modest chromatin differences between HFpEF and HFpEF-followup, identifying 94 DA peaks in type 2 and 63 in type 1 fibers (Fig 5c, Supplemental Dataset S3). Although few peaks passed the FDR threshold in the HFpEF-follow-up comparison, considering significant DA peaks in healthy control vs.

HFpEF comparison that were nominally significant (P<0.05) in HFpEF vs. follow-up revealed that the majority of peaks (94.7%) shifted in the opposite direction (Fig 5d). Example peak loci are shown in Fig 5e, and Extended Data Fig 7 which shows the *MAP2K6* gene, known to regulate muscle differentiation^107^, where shared non-DA peaks across clusters, along with a type 1 fiber and NMJ -specific DA peak, occur in the intronic region (black). Our analyses demonstrate that chromatin accessibility reverses towards the non-disease state post diet and exercise follow-up, similar to the RNA modality (Fig 4a).

To assess the chromatin context underlying the DA peaks, we computed enrichment for muscle fiber DA peaks in skeletal muscle ChromHMM states identified previously^108^. DA peaks between healthy controls and HFpEF in both type 1 and type 2 fibers were enriched in muscle active enhancer states and depleted in active TSS states (Fig 5f), suggesting a distal and more cell-type-specific regulatory function of DA peak elements.

Motif enrichment analyses of type 1 fiber DA peaks identified TFs involved in activity-dependent muscle metabolism (MEF2D, NR4A1)^109^, calcium signaling (NFATC4)^110,111^, and stress-responsive pathways (NR3C1, NFAT5)^112–114^, which, as in the gene expression modality, suggests a role of intrinsic skeletal muscle mechanisms such as metabolic efficiency, calcium pathways and stress adaptation in contributing to exercise intolerance in HFpEF (Fig 5g, Supplemental Table S9). Peaks showing reduced accessibility in HFpEF were preferentially enriched for nuclear receptor and endurance-related motifs, including MEF2D, NR4A1, NR1H2/3, and THRB^115^. In contrast, peaks with increased accessibility in HFpEF showed higher enrichment for stress-responsive and calcium-regulated pathway motifs, including AP-1 complex members (FOSB, JUND, ATF3)^116^, CREB family motifs^117^, and NFAT motifs. Notably, DA peaks that reversed following diet and exercise interventions were enriched for NFAT5 and MYF6 motifs, again implicating reversible activation of stress and remodeling programs in HFpEF skeletal muscle^118^.

Differential expression of TF genes was positively correlated with the enrichment of their cognate DNA-binding motifs in DA peaks (Fig 5h), indicating concordant changes in TF abundance and chromatin regulatory activity. TFs showing this coordinated regulation included regulators of oxidative stress and inflammation (BACH1, NFAT5)^112,119^, endocrine and metabolic signaling (THRB, ATF1, CREB1), and chromatin organization (NFYC, ZFHX4)^120,121^, highlighting a coherent and plastic regulatory architecture underlying skeletal muscle dysfunction in HFpEF.

Collectively, these results show that snATAC profiling identifies HFpEF-associated chromatin remodeling at muscle regulatory regions, which partially reverts towards the non-disease state following diet and exercise interventions.

### Integrating chromatin accessibility with gene expression reveals coordinated regulatory remodeling in HFpEF skeletal muscle

We leveraged our paired multiome data to understand the temporal dynamics between the epigenome and the transcriptome and how this relationship changed with HFpEF. We used MultiVeloVAE^122^, a Bayesian inference framework that models single cell/nucleus measurements of chromatin accessibility (c), unspliced (u), and spliced (s) RNA transcripts. A system of ordinary differential equations (ODEs) is used to infer latent variables including a unique cell time and cell state for each cell/nucleus, and for each gene in a cell infer a chromatin state (k_c_) and transcription rate (ρ) (Fig 6a). The decoder then reconstructs *c*, *u*, and *s* in cells using the ODE solutions. Inferred changes in *u*, *s*, and ρ values by condition (healthy control vs HFpEF) across genes correlated with observed gene expression fold changes (MAST), suggesting appropriate model fit (Extended Data Fig 8). The inferred velocity stream from our multiome data showed bifurcation at the mixed muscle fiber cluster level towards either type 1 or type 2 fiber cell types (Fig 6b). The multi-omic velocity inference approach also enabled characterizing how chromatin accessibility can influence gene transcription. Comparing cell- and gene-specific rate parameters k_c_ and ρ via the decoupling factor δ and coupling factor κ (Fig 6a) allows identifying scenarios where chromatin precedes transcription eg. chromatin opens before transcription begins (priming, decoupling factor δ = 1) or chromatin closes before transcription reduces (decoupling factor δ = -1); or when both modalities vary concordantly up or down (coupling factor κ = 1 or -1). Exploring when ATAC and RNA modalities were ‘out-of-sync’ via the decoupling factor δ, we found that genes showing increased priming between type 1 and type 1s cell types were enriched for chromatin regulator term (Fig 6c, Supplemental Table S11), including genes *MTF2*, *MBTD1* and *USP16* (Fig 6d-g Supplemental Table S10). Notably, the type 1s cluster was enriched in HFpEF nuclei (Fig 2e,f), which highlights the relevance of this priming signature in a fiber state that is over-represented in disease. Consistently, chromatin regulator genes were also enriched among genes with increased priming in HFPEF vs healthy control subjects (Kolmogorov-Smirnov test nominal p = 0.00011, not shown). Next, since most differentially expressed genes (60%) were downregulated in HFpEF (Fig 3b), we asked whether this shift was accompanied by a corresponding change in local chromatin dynamics. We found that more genes showed increased coupled repression in HFpEF, where chromatin accessibility and transcription both decline together (Fig 6h, Supplemental Table S12, binomial test P = 2.28e-6). These genes were enriched for skeletal muscle structural categories such as the sarcomere term (Fig 6i, Supplemental Table S13). The top ranked genes for this term included *CASQ1*, *FSD2*, and *JPH2*, which regulate the sarcoplasmic reticulum calcium storage and release -suggesting that repression of calcium-handling machinery in HFpEF is accompanied by coordinated closing of local chromatin (Fig 6j-l). Together, these analyses further highlight gene regulatory mechanisms of key chromatin regulators, and delineate the coordinated repression of the epigenome and transcriptome in processes related to calcium signaling in HFpEF skeletal muscle.

**Figure 6:**
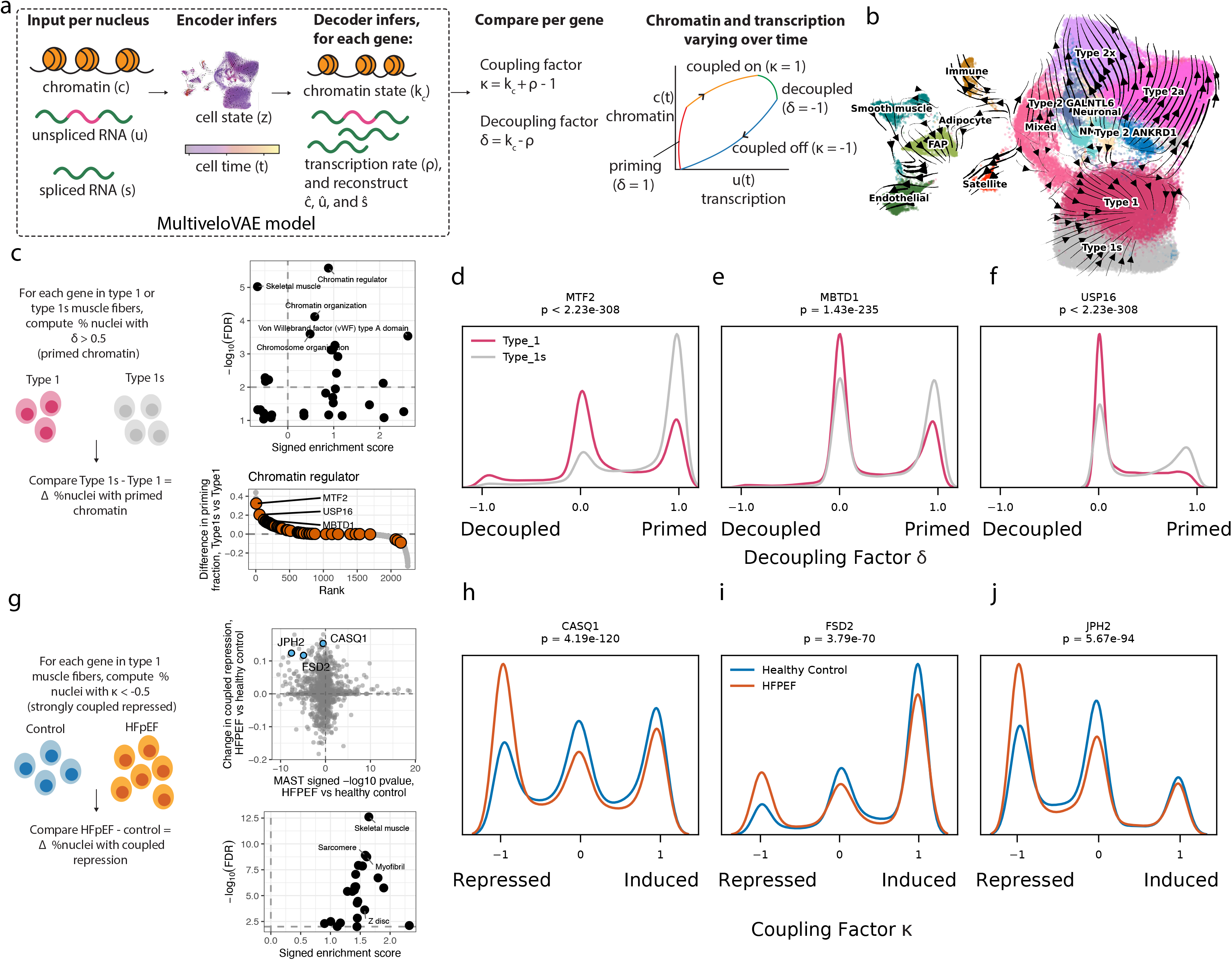
Multiomic velocity analyses identify temporal ordering between chromatin accessibility and transcription in HFpEF. a) MultiVeloVAE model estimates chromatin state k_c_ and transcription rate ρ for each gene in a nucleus, which through coupling and decoupling factors can indicate whether chromatin accessibility and transcription are co-ordinated (coupled-on or coupled-off) or decoupled. b) UMAP projection indicating the inferred velocity streams. c) Difference in chromatin priming for genes between type 1 and type 1s fibers was assessed by calculating fraction of nuclei with decoupling factor δ > 0.5. Volcano plot shows the functional enrichment for genes exhibiting increased priming in type1s vs type1 fibers. d) Rank plot of genes’ shift into priming in type1s relative to type1 fibers highlighting genes associated with Uniprot keyword “Chromatin regulator”. Distribution of decoupling factors δ for the chromatin regulators MTF2, MBTD1, and USP16 for type1 and type1s fibers are shown in e-g. (h) Difference in coupled repression for genes in type 1,1s fibers between control and HFpEF nuclei was assessed by calculating fraction of nuclei with coupling factor κ < -0.5. Scatter plot shows the change in fraction of cells in coupled-repression vs MAST differential expression signed -log p-value for each gene for healthy control vs HFpEF nuclei in type1,1s cluster. More genes showed downregulation in HFpEF and increased coupled repression (Binomial test P= 2.28e-6). (i) Volcano plot shows the functional enrichment for genes exhibiting increased coupled repression in healthy control vs HFpEF nuclei in type1,1s cluster. The coupling factor κ distributions in Type1/1s cells for sarcoplasmic reticulum and excitation-contraction coupling proteins CASQ1, FSD2 and JPH2 are shown in j-l for HFpEF and healthy control cells.

Having established relationships between chromatin accessibility and gene expression, we next sought to identify distal chromatin-gene links. Using a Poisson regression framework in SCENT (single-cell enhancer target gene mapping)^123^ we tested peaks where chromatin accessibility was associated with gene expression within 500kb of each DE gene TSS (Fig. 7a, Supplemental Dataset 4). We identified 4,198 gene-peak links for 910 DE genes in type 1 fibers (FDR < 10%, median 3 linked peaks per gene), and 3,324 gene-peak links for 517 DE genes in type 2 fibers (median 4 linked peaks per gene). Peaks closer to the TSS were more strongly linked to the gene (Fig 7b). Notably, peaks strongly linked to DE genes were more likely to be DA (odds ratio = 2.23, Fisher’s exact P 2.05e-24, Fig 7c), consistent with coordinated transcriptional and epigenomic remodeling.

**Figure 7:**
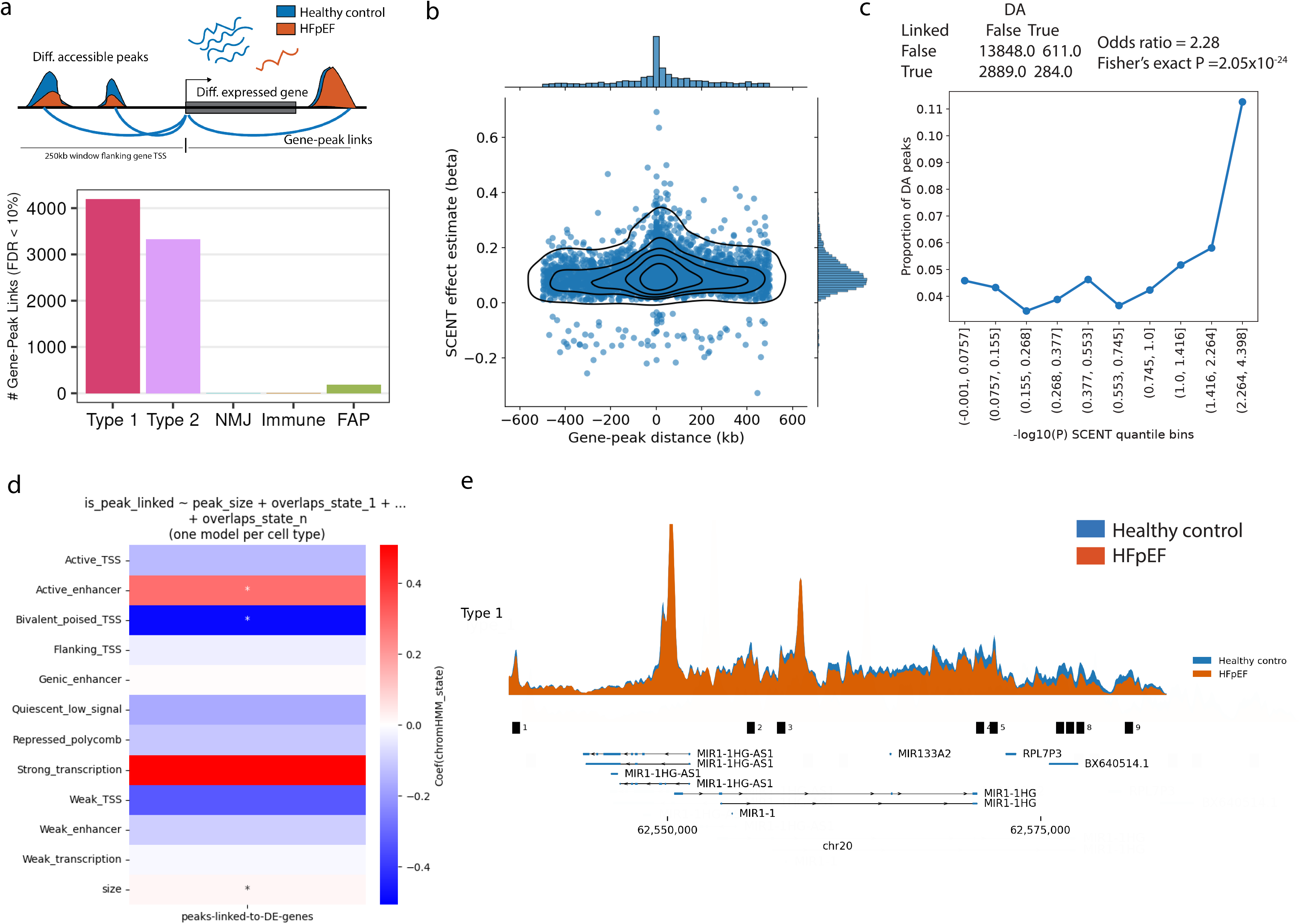
Multiome-based gene-peak linking identifies putative enhancers. a) Peaks within 500kb of TSS of DE genes were tested for chromatin accessibility association with gene expression using SCENT. Bars show the number of linked gene-peak pairs identified in each cluster (FDR < 10%). b) Gene-peak distance vs. SCENT effect size for linked peaks. c) Proportion of DA peaks by strata of linkage significance with DE genes. A 2x2 contingency table shows the number of peaks in type 1 fibers if these were identified as linked to a DE gene and DA by HFpEF. d) Enrichment for type 1 fiber peaks linked to DE genes in skeletal muscle chromatin states. e) Genomic locus of the *MIR1-1HG* gene showing 8 of the 49 identified linked peaks, which were differentially accessible by the HFpEF condition. *MIR1-1HG* is downregulated in HFpEF, and the DA peaks show reduced accessibility in HFpEF relative to healthy controls.

Peaks linked to DE genes in type 1 fibers were enriched in active enhancer chromatin states in skeletal muscle, further supporting a regulatory role of these identified putative enhancers (Fig 7d). In Fig 7e, we highlight the locus of the host gene for miR-1 (miR-1-1HG) - a long noncoding RNA regulator of myogenesis, fiber type specification, and mitochondrial function, known to target HDAC4, MEF2A/D, and PGC-1alpha pathways^124^. miR-1-1HG showed decreased expression in HFpEF muscle relative to healthy controls (log2(fold change) = -1.007, P = 1.34e-13). We identified 49 peaks where accessibility was significantly linked to miR-1-1HG expression; of these, 9 peaks were significantly differentially accessible, 8 of which exhibited reduced accessibility in HFpEF. These results delineate broad chromatin remodeling at a central myogenic locus, highlighting the link between enhancer accessibility and gene expression changes at key regulators of skeletal muscle function in HFpEF.

### Integration with cardiometabolic GWAS signals helps nominate disease mechanisms

We next asked if our identified regulatory elements were enriched to overlap with complex trait and disease-associated genetic variation. snATAC peaks in muscle fibers were enriched in GWAS signals for diastolic blood pressure, pulse rate, heart failure, and atrial fibrillation (Fig 8a, Supplemental Table S14), indicating that genetic variants influencing these cardiovascular traits are preferentially enriched in skeletal muscle regulatory elements. Skeletal muscle fiber peaks were also enriched in type 2 diabetes and waist-to-hip ratio (WHR) (both adjusted for BMI) signals, indicating their importance in metabolic pathways. Notably, DA peaks in type 1 fibers exhibited higher fold enrichment estimates in both cardiovascular and metabolic traits than all peaks in clusters. Moreover, DA peaks linked to DE genes in type 1 were enriched for insulin fold change, a measure of insulin resistance in peripheral tissues, and WHR, which suggests that our DA and gene-peak linking analyses identified disease-relevant metabolic regulatory signatures.

**Figure 8:**
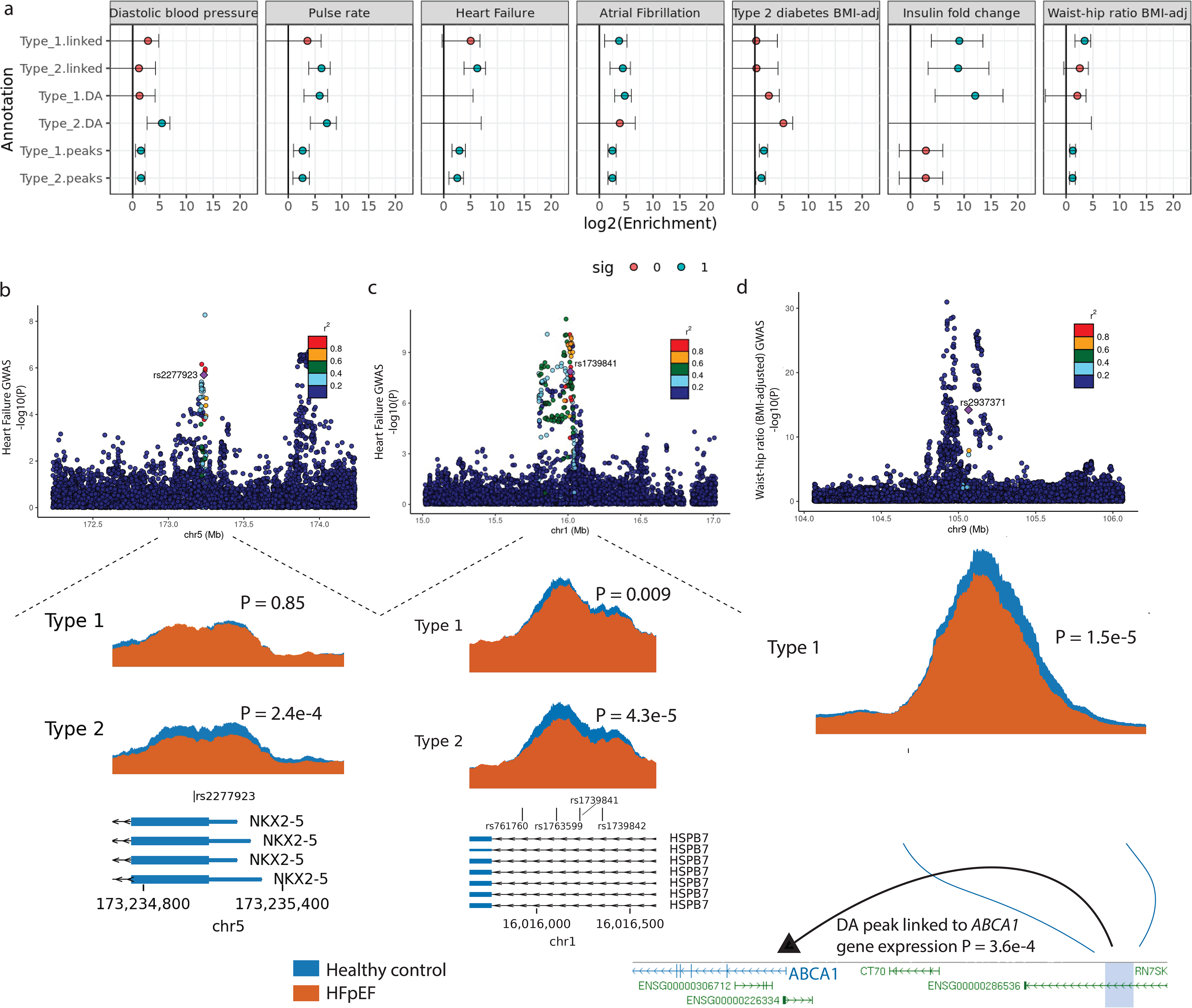
Integration with GWAS signals nominates causal variants and target genes. a) Enrichment for cluster snATAC peaks, muscle fiber DA peaks, and peaks linked to DE genes in HFpEF to overlap cardiometabolic GWAS signals. Example GWAS signals where a 95% credible set variant(s) overlap HFpEF DA peaks, along with orthogonal data such as gene-peak linking, peak-peak linking, and eQTL information nominating effector transcripts. Shown are heart failure GWAS signals at the *NKX2-5* locus (b), and the *HSPB7* locus (c), along with a waist-hip ratio adjusted for BMI GWAS signal at an intergenic locus on chr9 that was linked to the *ABCA1* gene expression (d).

We overlapped DA peaks with GWAS signal 95% credible set variants to examine if integrating chromatin accessibility could help prioritize likely causal variants, and combined our gene-peak links, snATAC peak-peak coaccessibility (Cicero^125^), along with publicly available eQTL information to nominate target genes. At a genome-wide-significant heart failure GWAS signal on chr5, one credible set variant rs2277923 (GWAS P = 2e-6, PIP=0.03) overlaps a type 2 fiber-specific DA peak in the promoter region of the *NKX2-5* gene (DA log2(FC) = -0.412, P = 2.4e-4, Fig. 8b). Risk allele T at this variant is associated with increased *NKX2-5* gene expression level in GTEx pituitary gland tissue (normalized effect size, NES = 0.86, P = 6.3e-34)and was a splice-QTL for NKX2-5 in heart tissue (NES = 0.3, P = 2.9e-8)^126^. NKX2-5 is a key developmental TF that has also been implicated in skeletal muscle biology, where its overexpression in mouse myoblasts was shown to inhibit myogenic differentiation and regeneration^127^. These data nominate *NKX2-5* as a candidate effector gene at this heart failure risk locus through regulatory disruption in type 2 fibers.

At another heart failure GWAS signal on chr1, four variants in high linkage disequilibrium (LD) with the index SNP rs1763604 (r2 > 0.91 with the index SNP, GWAS P < 1.5e-8) overlapped a DA peak in the *HSPB7* gene intron in type 2 fibers (log2(FC) = -0.255, P = 4.3e-5, Fig 8c).

Peak-peak co-accessibility analyses with Cicero identified that the DA peak was co-accessible with the *HSPB7* promoter peaks, implicating the gene as a likely target. *HSPB7* encodes a heat-shock protein CVHSP, which is primarily expressed in the heart, and acts as a chaperone to maintain sarcomeric integrity and structure^128,129^.

At a waist-hip ratio adjusted for BMI (WHR-adjBMI) GWAS locus on chr9, among independent GWAS signals, SNP rs2937371 tagged a secondary, yet highly significant signal (P=6.031e-15, credible set PIP=0.99) and overlapped a DA peak in type 1 fibers. The overlapping peak was linked to the expression of the *ABCA1* gene ∼139kb away (Fig. 8d). The peak exhibited reduced accessibility in HFpEF (log2(FC) = -0.24, P = 1.5e-5), and *ABCA1* expression was concordantly downregulated in HFpEF (log2(FC) = -0.7, P = 4.0e-6). Reduced *ABCA1* expression in muscle in the mouse has been shown to increase intramyocellular cholesterol accumulation, impair insulin-dependent AKT phosphorylation and GLUT4 translocation, thereby accelerating insulin resistance^102^. While downregulated in HFpEF, *ABCA1* expression is restored towards baseline after diet and exercise followup (log2(FC)=0.49, P=1.1e-13). Together, these findings link a WHR-adjBMI genetic risk locus to HFpEF-associated chromatin remodeling and altered metabolic gene regulation in skeletal muscle.

## Discussion

We performed single-nucleus multiomic profiling of 132,568 high-quality nuclei isolated from 140 *vastus lateralis* skeletal muscle biopsies from 95 participants, including healthy age-matched controls, HFpEF patients at baseline, and HFpEF patients following a supervised diet and exercise intervention. This is the first study of its kind to profile both gene expression and chromatin accessibility in the skeletal muscle of human HFpEF patients. As a major organ in metabolic dysfunction and exercise intolerance in HFpEF, understanding the molecular dynamics of HFpEF skeletal muscle is essential to a fuller understanding of the disease and potential treatments. Additionally, paired profiling of patients before and after diet and exercise enabled understanding the mechanisms behind how these interventions alleviate exercise intolerance. A key strength of our study is its randomized block design, where biopsies were pooled into covariate-balanced batches and resolved by genotype-based demultiplexing; which guarded our experimental contrasts against technical batch effects, enabled genetic doublet detection, reduced costs, and overall increased the rigor of the work.

Our analyses revealed disease-associated remodeling of skeletal muscle composition and transcriptional programs. We identified a distinct HFpEF-enriched population of type 1 muscle fibers, termed type 1s, which were transcriptionally recognizable as slow-twitch fibers but exhibited dysregulated CAMK2D signaling, including increased CAMK2D and decreased NFAT-calcineurin pathway activity. Chronic CAMK2D activation in skeletal muscle has been associated with muscle atrophy, while suppression of NFAT-calcineurin signaling is known to impair endurance-related transcriptional programs, promoting a transition from type 1 fibers to more easily exhaustible type 2x fibers^130,131^. Across major muscle fiber groups, we observed disruptions in metabolic processes and cytoskeletal structure. These changes are consistent with a transition toward a metabolically less efficient, type 2x–like phenotype, which may contribute to impaired endurance and exercise intolerance. Differential abundance analyses also confirmed enrichment of type 2x fibers and depletion of canonical type 1 fibers and pericytes in HFpEF, reinforcing the contribution of both fiber and vascular remodeling to skeletal muscle dysfunction.

The predominance of downregulated genes in HFpEF relative to healthy controls (∼60%), points to a loss-of-function signature as the primary driver of impaired muscle endurance. The impairments we observe in skeletal muscle share overlap with pathways contributing to heart disease, as indicated in the enrichment for the hypertrophic cardiomyopathy KEGG term. This convergence suggests a shared disease origin, in which systemic transcriptional dysregulation driving cardiac hypertrophy simultaneously induces an atrophic response in peripheral muscle. The combined dysregulation of muscle cytoskeleton components alongside impaired oxytocin and calcium signalling implies that HFpEF muscle enters a maladaptive remodeling state. These findings agree with the framing of HFpEF as a systemic disease with coordinated, cross-tissue consequences.

We demonstrated that HFpEF is associated with remodeling of the skeletal muscle regulatory landscape, marked by coordinated changes in chromatin accessibility, TF motif enrichment and gene expression. Differentially accessible peaks were enriched in active enhancer states and showed strong concordance with differential gene expression, supporting a direct regulatory role for these elements. Directional motif analyses revealed a shift from accessibility at nuclear receptor and endurance-associated TF motifs (including MEF2D, NR4A1, NR1H2/3, and THRB) toward increased accessibility at stress-responsive and calcium-regulated motifs such as AP-1 complex members, CREB, and NFAT family TFs. This pattern is consistent with suppression of oxidative and metabolic programs alongside activation of stress and remodeling pathways, aligning with peripheral mechanisms of exercise intolerance in HFpEF.

At the level of both gene expression and chromatin accessibility, we observed striking reversibility of HFpEF-associated alterations following diet and exercise. MAST-based analyses demonstrated that a majority (∼87%) of HFpEF DE genes across skeletal muscle fiber types reversed toward the healthy state after intervention. Reversing genes in type 1 fibers were enriched in metabolic and cytoskeletal pathways, including restoration of mitochondrial and fatty acid oxidation genes alongside normalization of actin and cytoskeletal organization.

Epigenomically, chromatin accessibility also trended towards the non-disease state at nuclear receptor and oxidative metabolism-associated regulatory elements. Together, these findings highlight the plasticity of skeletal muscle regulatory programs in HFpEF and support a model in which impaired metabolic and activity-dependent signaling contributes to exercise intolerance, while lifestyle interventions can partially restore these pathways.

Paired chromatin accessibility and transcription profiles in the same nuclei allowed us to move beyond static comparisons and infer the temporal ordering of epigenetic and transcriptional changes. The inferred velocity stream positioned the mixed muscle fiber cluster at a bifurcation point from which trajectories emanated toward type 1 or type 2 fiber identities. Nuclei in this cluster expressed markers for both canonical fiber types, which is consistent with these being hybrid or transitional myofibers. The mixed fiber state is a known feature of skeletal muscle, having been observed in previous snRNA studies, especially during the processes of aging and regeneration^132–136^.

Considering cell- and gene-specific rate parameters from the multi-omic velocity analyses, two key patterns emerged. First, chromatin remodelers, including *MTF2*, *MBTD1*, and *USP16*, showed evidence of increased chromatin priming -where accessibility increases ahead of transcription -in the HFpEF-enriched type 1s cluster, suggesting that chromatin opening at key regulatory factors is relevant in disease-related fiber type identity. Second, and consistent with the predominance of downregulated genes in HFpEF, we found an overall skew in genes toward coupled repression, in which loss of accessibility and loss of transcription occur together. This coupling was pronounced at calcium-handling genes critical for sarcomere function such as *CASQ1*, *FSD2*, and *JPH2*, indicating that the impaired calcium signaling characteristic of HFpEF muscle is not solely a transcriptional phenomenon but is reinforced at the level of local chromatin.

Beyond disease-associated remodeling, HFpEF-linked differentially accessible peaks were significantly enriched for diastolic blood pressure, pulse rate, and atrial fibrillation GWAS signals-further demonstrating the shared nature of cardiac pathways in skeletal muscle regulation.

Furthermore, peaks linked to DE genes in HFpEF were enriched for insulin resistance and waist-to-hip ratio GWAS signals, indicating that transcriptional reprogramming in HFpEF skeletal muscle is linked to regulatory elements that are genetically linked to insulin resistance and adiposity, supporting that obesity-related HFpEF is a systemic, metabolic disorder involving substantial skeletal muscle abnormalities that contribute to symptoms and impaired exercise capacity, in addition to cardiac abnormalities.

Integration of GWAS 95% credible set variants with chromatin accessibility, gene–peak linking, co-accessibility, and eQTL data nominated causal variants and effector genes, providing mechanistic links from genetic risk to regulatory disruption and skeletal muscle dysfunction. At a heart failure locus on chr5, we nominated *NKX2-5,* which encodes a developmental transcription factor that restrains myogenic differentiation and regeneration, suggesting a route by which this risk allele could impair muscle repair. At another heart failure locus on chr1, integrative analyses implicated *HSPB7,* which encodes the cardiac chaperone CVHSP that maintains sarcomeric integrity. These findings suggest that heart failure-associated variation may influence fiber-type-specific chromatin remodeling and engage shared stress-response and structural maintenance pathways across skeletal and cardiac muscle. At a WHR-adjBMI signal on chr9, reduced accessibility of a type 1 fiber peak in HFpEF linked to concordantly reduced expression of *ABCA1*, a regulator of intramyocellular cholesterol and insulin signaling. Notably, *ABCA1* expression was partially restored towards baseline following diet and exercise, implicating a reversible, intervention-responsive node linking WHR-related genetic risk to HFpEF-associated metabolic dysfunction in muscle.

Our analyses were most robust in the abundant muscle fiber populations, and lower-abundance cell populations were underpowered for comprehensive regulatory analysis. Nevertheless, the high quality of the data, paired multiome profiling of HFpEF patients enabled detection of disease-associated and likely intervention-responsive regulatory signatures in specific muscle fibers.

In summary, HFpEF is associated with skeletal muscle remodeling, encompassing type 1 fiber subtype transitions, vascular and FAP alterations, and dysregulated calcium-mediated and metabolic signaling. These molecular signatures are partially reversible at both the transcriptional and epigenomic levels following lifestyle intervention, emphasizing the therapeutic potential of targeting skeletal muscle in addition to the heart in HFpEF. This study provides a comprehensive resource linking cellular composition, transcription, chromatin accessibility, and genetics, illuminating the peripheral mechanisms that contribute to exercise intolerance in HFpEF.

## Acknowledgements

We would like to thank the Parker lab members for their help and feedback for this work. This work was supported by grant Collaborative Research Agreement 673400 from Pfizer Inc.

## Author Contributions

MB, AV, NC performed data processing and computational analyses, interpreted and visualized the results, prepared figures and wrote the manuscript. PO performed data preprocessing. MP, EP provided analyses insights. VD performed the multiome experiments. NM, EC performed genotyping. DK, BN, and PB conducted the SECRET I and II trials and the Health Control Study which provided the biological samples, and provided content expertise and reviewed and edited the manuscript. RJRF, MP, EP, EBF, and FD provided intellectual contribution to the analysis workflow and interpretation of results, and provided funding. SCJP organized the project and supervised all aspects of data generation, analyses and interpretation, and edited the manuscript.

## Methods

### Muscle biopsy sample collection

Frozen needle biopsy samples from the *vastus lateralis* muscle of participants of the SECRET I^28^ and SECRET II^25^ randomized, controlled trials and healthy control cohort, were obtained according to previous published techniques by the team at Wake Forest University School of Medicine^15^. Participants from SECRET I were assigned to one of four 20-week intervention conditions: 1) a supervised hypocaloric diet prepared by the Wake Forest Clinical Research Metabolic Kitchen under the direction of a registered dietitian, 2) a supervised exercise condition consisting of 1 hour of supervised walking with progressive intensity, 3 times per week, 3) a combined diet and exercise intervention, or 4) no intervention. Participants from SECRET II all underwent a supervised hypocaloric diet in the same manner as SECRET I participants.

Samples were collected before and after a 20-week supervised exercise condition consisting of 3 hour-long sessions of either aerobic training, or aerobic training with additional resistance training.

### Batch assignment

We processed muscle biopsy samples in 7 batches of 20. Biopsies were assigned to batches using OSAT^137^. We used this tool to distribute the 140 total biopsies across 7 multiome batches, such that the average age, bmi, and biological sex distribution was approximately the same across batches. The batches were also matched for disease status and study. Baseline and follow-up biopsies from the same SECRET II participant were deliberately assigned to different batches, ensuring that every sample within a pooled batch derived from a distinct individual and could therefore be resolved by genotype-based demultiplexing.

### Nuclei isolation and sorting

Frozen biopsies were cryopulverized under liquid nitrogen. For each batch, pulverized tissue from the 20 constituent biopsies was combined as four pools of five samples, yielding approximately 200 mg of tissue per batch. All buffers were prepared fresh, all steps were performed on ice and all centrifugation was carried out at 4 °C. Each of the four pools was resuspended in 1 mL of lysis buffer (50 mM HEPES pH 7.5, 140 mM NaCl, 1 mM EDTA pH 8.0, 10% glycerol, 0.5% NP-40, 0.25% Triton X-100, 5 µg/mL 7-AAD (BioLegend, 420404), 1 U/µL Protector RNase inhibitor (Roche, 3335399001) and 1 tablet EDTA-free cOmplete mini protease inhibitor (Roche, 11836170001)), and the four suspensions were then combined in a 7 mL Dounce homogenizer to give a single 4 mL lysate per batch.

Tissue was dispersed by gentle pipetting until no macroscopic fragments remained and then homogenized with 15 strokes of the loose (A) pestle. The homogenate was divided between two 2 mL tubes and lysed further on ice by pipetting with a wide-bore tip at 9 strokes per minute for 5 min. Nuclei were pelleted at 500g for 5 min, resuspended in 1 mL of ice-cold wash buffer (10 mM Tris-HCl pH 7.4, 10 mM NaCl, 3 mM MgCl₂, 1% BSA, 0.1% Tween-20, 1 mM DTT, 5 µg/mL 7-AAD and 1 U/µL Protector RNase inhibitor) and centrifuged at 100g for 1 min to sediment residual debris. The nuclei-containing supernatant was retained and passed sequentially through duplicate 70 µm, 50 µm and 30 µm strainers, each pre-wet with 200 µL of 1% BSA and chased with 200 µL of wash buffer after the sample had passed. Filtered nuclei were pelleted at 500g for 5 min and resuspended in 3 mL of sort buffer (1% BSA, 0.1% Tween-20, 1 mM DTT, 5 µg/mL 7-AAD and 1 U/µL Protector RNase inhibitor in Ca²⁺- and Mg²⁺-free PBS).

After a 15 min incubation on ice to allow 7-AAD labelling of nuclear DNA, the suspension was filtered through a 20 µm strainer and 7-AAD-positive events were purified on a MACSQuant Tyto (Miltenyi Biotec). This microchip-based sorter operates at low pressure and avoids the shear forces of conventional sorting, which we found helped with preserving nuclear integrity in this tissue. Sorted nuclei were counted on disposable hemocytometer slides and assessed by light microscopy to confirm intact, well-separated nuclei free of visible debris and membrane blebbing.

### Library preparation and sequencing

Purified nuclei were submitted to the Advanced Genomics Core at the University of Michigan for single-nucleus multiome profiling (paired 3′ snRNA-seq and snATAC-seq) on the 10x Genomics Chromium platform using the Chromium Next GEM Single Cell Multiome ATAC + Gene Expression kit. Additional 5′ snRNA-seq was performed on 60 muscle biopsies as part of a separate ongoing project. Light whole-genome sequencing of the same participants was performed to enable downstream genetic demultiplexing of each batch.

### Genotyping and imputation

We performed light whole-genome sequencing on the 95 individuals. The DNA was extracted from the muscle tissue using qiagen kit DNeasy Blood & Tissue Kit (50) Cat no. / ID. 69504 and submitted to the University of Michigan Advanced Genomics Core for light whole-genome genome sequencing.

We performed genotype imputation using GLIMPSE-2^138^ with 1000G as the reference panel. We downloaded the 1000G reference from http://ftp.1000genomes.ebi.ac.uk/vol1/ftp/data_collections/1000G_2504_high_coverage/working/20201028_3202_phased/CCDG_14151_B01_GRM_WGS_2020-08-05_chr${chrom}.filtered.shapeit2-duohmm-phased.vcf.{gz,gz.tbi}, selected only bi-allelic variants and converted to the bcf format using bcftools. We split the genome into chunks define to run imputation and phasing using GLIMPSE2_chunk. We then converted the reference panel into GLIMPSE’s binary format using GLIMPSE2_split_reference and performed imputation and phasing using GLIMPSE2_phase. We ligated the imputed chunks using GLIMPSE2_ligate.

### snATAC-seq data processing

Single-nucleus ATAC-seq data were processed using the Parker Lab snATAC-seq pipeline, available at https://github.com/porchard/snATACseq-NextFlow (commit 89ab7d5). We trimmed sequencing adapters using cta (v. 0.1.2) and mapped reads to hg38 with bwa (v.0.7.15; bwa mem -I 200,200,5000 -M)^139^. Cell barcodes were corrected using a custom python script implementing the barcode correction algorithm described on the 10x Genomics website.

Duplicates were marked using picardtools (http://broadinstitute.github.io/picard/; v 2.26.9; MarkDuplicates READ_ONE_BARCODE_TAG=CB READ_TWO_BARCODE_TAG=CB VALIDATION_STRINGENCY=LENIENT). BAM files were filtered to properly mapped, non-duplicate autosomal read pairs using samtools (samtools view -h -b -f 3 -F 4 -F 8 -F 256 -F 1024 -F 2048 -q 30 chr{1..22}). QC metrics were generated using ataqv (v. 1.3.0;--ignore-read-groups --nucleus-barcode-tag CB, with the blacklist from ENCODE (https://www.encodeproject.org/files/ENCFF356LFX/))^140^. Code for these steps can be found at https://github.com/porchard/snATACseq-NextFlow (commit 89ab7d5)

### snRNA-seq data processing

Single-nucleus RNA-seq data were processed using the Parker Lab snRNA-seq pipeline, available at https://github.com/porchard/snRNAseq-GitHub-porchard/snRNAseq-NextFlowNextFlow. Reads were mapped to the hg38 genome using STARsolo v.2.7.10a and filtered to exclude secondary alignments, supplemental alignments, and alignments with mapping quality less than 255 with samtools^141,142^. Ambient contamination correction was performed with Cellbender^143^. Empty droplets and low-quality nuclei were identified using dropkick. Ambient contamination was removed using Cellbender.

### Nuclei QC

After multiome snRNA and snATAC preprocessing, we applied several QC thresholds on both RNA and ATAC barcodes. For RNA barcodes, we considered: minimum UMI = 120, maximum mitochondrial read fraction = 0.1, minimum probability of singlet form cellbender = 0.99, post-cellbender UMIs = 100, maximum fraction of exon vs full-gene-body reads = 0.6. For ATAC barcodes, we considered library-specific minimum high-quality autosomal alignments (HQAA) thresholds, which ranged from 3000 to 10000, and minimum TSS enrichment = 2.

### Demultiplexing, doublet detection, and nuclei filtering

We used Demuxlet^144^ (v. 2, popscle package https://github.com/statgen/popscle) to identify high-quality droplets containing single nuclei (“singlets”) and assign them to samples based on genotypes. For snRNA, we considered variants in gene bodies (Gencode v30^145^); for snATAC, we considered variants in peaks identified across skeletal muscle cell types from a previous muscle snATAC study^132^, since these regions concentrate the majority of informative reads and minimize background noise. We ran Demuxlet with the snRNA/snATAC library bam files and genotype vcf files containing samples included in the respective batch, using default parameters and selected singlet droplets.

### Clustering and cell type annotation

RNA nuclei were clustered using Seurat v5^29^. This analysis included 102,941 multiome singlet nuclei that passed all RNA and ATAC QC thresholds, and 29,627 multiome singlet nuclei that passed all RNA QC thresholds only. In addition, 61,565 5’ gene expression (GEX) nuclei were included in the clustering as part of ongoing related projects. In Seurat, we used the sctransform approach for normalization and variance stabilization while adjusting RNA mitochondrial fraction and exon to full-gene-body read fractions (SCTransform function). We then computed principal components (RunPCA) and integrated layers using the Harmony integration method (IntegrateLayers function). Next, we ran the Seurat functions FindNeighbors with 25 harmony dimensions and FindClusters with the Louvain algorithm and resolution = 0.3. This process resulted in 15 distinct cell-type clusters which we annotated using known marker gene expression. We performed subclustering on the smooth muscle, endothelial, FAP, and immune cell nuclei by separating these nuclei and running FindNeighbors and FindClusters over a range of Harmony reduction dimensions and Louvain clustering resolutions, and selected the subclustering that provided biologically meaningful sub-clusters such as identifying pericytes from smooth muscle; macrophage and T cell from immune cells; capillary, arterial, venous, and lymphatic subtypes from endothelial cells; and progenitor, fibrogenic, and adipogenic subtypes from FAPs.

### Differential gene expression analysis

Differential gene expression analysis was performed using MAST^51^. For each cell type, we tested for 1) association of gene expression with disease (relative to healthy samples) and2) association of gene expression with diet+exercise treatment (in HFpEF samples, relative to samples from the same subjects before diet and exercise treatment). We created filtered gene lists for the two contrasts consisting of genes expressed in at least 10% of cells for the samples included in the contrast. We then used the union of the two lists as the final set of genes to be tested for that cell type. This allowed us to compare results between contrasts.

Genes counts were normalized by library size using transcripts-per-million and then log2-transformed with a pseudocount. We modeled gene expression for each cell type using MAST’s zlm function with method=”glmer”, ebayes=FALSE, strictConvergence=FALSE, and fitArgsD=list(nAGQ = 0). strictConvergence=FALSE is a parameter recommended by the authors of MAST to account for the underlying glmer function’s sensitivity to warnings when running on large datasets. fitArgsD=list(nAGQ = 0) is a parameter recommended by the authors to improve the speed of running the model (from multiple weeks to within a day, for the largest cell type group in our data) by using a faster but less exact form of parameter estimation.

For testing the association of gene expression with disease, we used age, batch, sex, and the cellular detection rate (calculated as the number of genes detected in a sample) as fixed-effect covariates, and sample ID as a random effect covariate. Because BMI distributions were significantly different between healthy controls and HFpEF participants (Extended Data Fig 9), we could not control for BMI without controlling for the effect of disease overall. For testing association of gene expression with diet+exercise treatment, which utilizes paired samples, we used batch, subject ID, and the cellular detection rate (calculated as the number of genes detected in a sample) as fixed-effect covariates, and sample ID as a random effect covariate.

For each model, we performed the likelihood ratio test (LRT) to test for association between gene expression and 1) disease status (relative to healthy controls) or 2) diet+exercise treatment (in HFpEF samples, relative to samples from the same subjects before diet and exercise treatment). Multiple-testing correction was performed for all results using the Benjamini-Hochberg procedure. One individual enrolled in both SECRETI and SECRETII and provided one biopsy in SECRETI and paired biopsies in SECRETII. To keep the independence assumption of biopsies the control vs HFpEF comparison, we removed the SECRETI biopsy (sample id 6721-NM-56) from this analysis since it had fewer nuclei.

### Differential abundance analysis

We computed a k-nearest neighbour graph of our nuclei using miloR’s buildGraph function with parameters k=70, d = 25^30^. Then, vertices were grouped into neighborhoods using makeNhoods (prop = 0.05, k = 7-, d = 25), and the number of nuclei per sample in each neighborhood were counted using countCells. Neighborhoods were annotated by their majority cell type, according to established nuclei annotations. We tested for differential abundance of HFpEF nuclei relative to No-HFpEF nuclei across all neighborhoods using the testNhoods function, with age, batch, and sex as covariates. We used the function findNhoodGroupMarkers to test for differentially expressed marker genes between type 1s neighborhoods with significant (spatial FDR < 5%, which accounts for overlapping nuclei between neighborhoods) differential abundance and a positive log2FC (enriched in HFpEF) and type 1 neighborhoods with significant differential abundance and a negative log2FC (enriched in healthy). Gene set enrichment analysis on marker genes reaching BH-corrected p-value < 0.01 was performed as described below.

### Gene set enrichment analysis

The enrichGO and enrichKEGG functions from clusterProfiler were used to perform gene set enrichment analysis on all nominally significant (pvalue < 0.05) genes for each celltype, for each contrast^146^. We tested for gene sets from the following databases: Kyoto Encyclopedia of Genes and Genomes (KEGG) pathways, Gene Ontology (GO) biological processes, cellular components, and molecular functions. The statistical background for each test was restricted to the set of input genes used to run MAST, which were obtained as described above. Pathway p-values were corrected using the Benjamini-Hochberg procedure. GeneRatio was calculated as the ratio of nominally significant genes to the number of genes in the pathway when restricted to genes in the background set.

### snATAC peak calling

We processed the snATAC data for 102,941 pass-QC multiome nuclei in SnapATAC2^147^(v 2.7.1). We mapped the RNA-clustering annotations and called peaks within clusters using macs3 within SnapATAC2, considering reproducible peaks across batches (providing replicate = “batch” in tl.macs3 function and replicate_qvalue = 0.05). We then used the tl.merge_peaks function to call consensus peaks across clusters. Peak summits are initially expanded by half_width = 250bp on both sides. Following this expansion, it addresses the issue of overlapping peaks through an iterative process. The procedure begins by prioritizing the most significant peak, determined by the smallest p-value. This peak is retained, and any peak that overlaps with it is excluded. Subsequently, the same method is applied to the next most significant peak. This iteration continues until all peaks have been evaluated, resulting in a final list of non-overlapping peaks, each with a fixed width determined by the initial extension.

### Differential chromatin accessibility analysis

We tested peaks for differential accessibility by disease condition in clusters using a nucleus-level negative-binomial generalized linear mixed model (NB-GLMM) approach. For the control vs HFpEF comparison, we modeled peak counts ∼ disease condition + age + sex + batch + offset(log(total_counts)) + (1|sample), where the biopsy sample id is taken as a random effect to account for multiple nuclei from a single sample, and log(total_counts) is taken as an offset. We then fit a null model keeping all covariates but without the disease condition term, and performed a likelihood ratio test (LRT) to test if including the disease condition term increased the likelihood of the model indicating an association with disease. As described for the differential gene expression analysis methods, we removed sample id 6721-NM-56 from this analysis.

For the HFpEF vs HFpEF-followup comparison where we had matched biopsies, we included individual as a fixed effect to account for within-individual pairing, and sample was included as a random effect to account for multiple nuclei per biopsy. Age and sex were not included in this model as they were constant within individuals. peak counts ∼ disease condition + batch + individual + offset(log(total_counts)) + (1|sample)

### Multi-omic Velocity Analysis

RNA reads were aligned using starsolo as described above (snRNA-seq data processing) with the Velocyto output option specified. Nuclei with multiome data that passed the QC thresholds indicated above (Nuclei QC, 102,941 nuclei) were loaded into an annData object (scanpy v 1.12)^148^. RNA counts were normalized per cell, subset to the 3000 variable genes used above, log transformed and scaled. Unspliced and spliced moments were calculated using 30 PCs and 30 nearest neighbors. ATAC peaks were generated by SnapATAC2 as described above (snATAC peak calling). Peaks were linked to genes as a promoter if they overlapped a region within 1000bp of its TSS, as distal peaks if they were within 100kb of its TSS or overlapped the transcript body or as intergenic peaks if they were correlated with gene expression as determined by Signac (p < 0.05). Peak-gene links were calculated for each coarse nuclei cluster. Counts for linked peaks were summed for each gene to generate a single ATAC value for each nuclei for each gene. ATAC summed counts were TF-IDF normalized and smoothed across their 50 nearest neighbors. Nearest neighbors were calculated from RNA and ATAC data using the Seurat FindMultiModalNeighbors function.

RNA and ATAC data were submitted to MultiVeloVAE (v0.1.0)^122^ for velocity modeling conditioned on batch and regressing out unspliced rna count. Splicing and chromatin velocities were coupled-increasing if their coupling factor κ was > 0.5 and coupled-decreasing if k < -0.5. They were decoupled if their decoupling factor δ was > 0.5 or < -0.5. For each gene the fraction of cells that were in each state was calculated and compared between disease states or between cell clusters. The difference in the coupled-repressed fraction between subjects with HFpEF and healthy controls was calculated for each gene and submitted for rank-based functional enrichment analysis using a Komogorov-Smirnov test as implemented in the String Database functional enrichment tool (https://string-db.org/)^149^. A signed enrichment score was calculated for each category as the W statistic multiplied by the direction of the shift (-1 for top of the list, 1 for bottom)

### Analysis of gene-peak links

To associate gene expression with chromatin accessibility, we used SCENT^123^ to test gene-peak links within clusters. For each DE gene and a peak within 500kb of its TSS, we fit Poisson regression models relating gene expression counts over binarized peak accessibility. We included covariates for age, sex, batch, disease group (healthy control, HFpEF or HFpEF-followup), snRNA mitochondrial fraction, nucleus-level log-transformed total UMI counts, and snATAC TSS enrichment. Biopsy-level sample identity was not included due to collinearity with batch and disease group. To improve computational efficiency, we modified the SCENT implementation to use the fastglm library (https://github.com/jaredhuling/fastglm) in place of glm. SCENT performs iterative bootstrapping to estimate beta coefficients and a two-sided p value. We considered gene-peak links at FDR < 10% using Storey’s qvalue approach.

### Peak co-accessibility

We ran Cicero^125^ (v. 1.4.0) on the peak fragment counts in each cluster to score peak-peak co-accessibility. We processed snATAC data in Signac, to derive a low-dimensional embedding for use as the manifold input to Cicero. Peaks were TF–IDF normalized (RunTFIDF) and all peaks retained as features (FindTopFeatures, min.cutoff = ’q0’). Latent semantic indexing (LSI) was performed by singular value decomposition (RunSVD), and a UMAP embedding was computed from LSI components 2–50 (RunUMAP), excluding the first LSI component, which is typically correlated with sequencing depth rather than biological signal. This ATAC LSI UMAP embedding was then used as the reduced coordinates and set window size to 500 kb. A peak was considered to be a TSS peak for a gene if it overlapped the 1kb window upstream of that gene’s TSS. If multiple TSS peaks were present for a gene, the maximum co-accessibility score was considered.

### GWAS enrichment

We computed enrichment for each peak annotation to overlap GWAS signals using fgwas^150^ (v. 0.3.6) using default parameters. We processed GWAS summary statistics as required by the program, using liftOver to lift to hg38 when required, and provided columns CHR POS F Z N SE SNPID.

## Data Availability

We are in the process of making the raw data available through the European Genome-phenome Archive (EGA). Processed data is submitted to Zenodo (DOI: 10.5281/zenodo.22083225).

### Code Availability

The code used to run analyses in this work are available on GitHub (https://github.com/ParkerLab/2026_HFpEF_multiome_mbose).

---

**Extended Data Fig 1:**
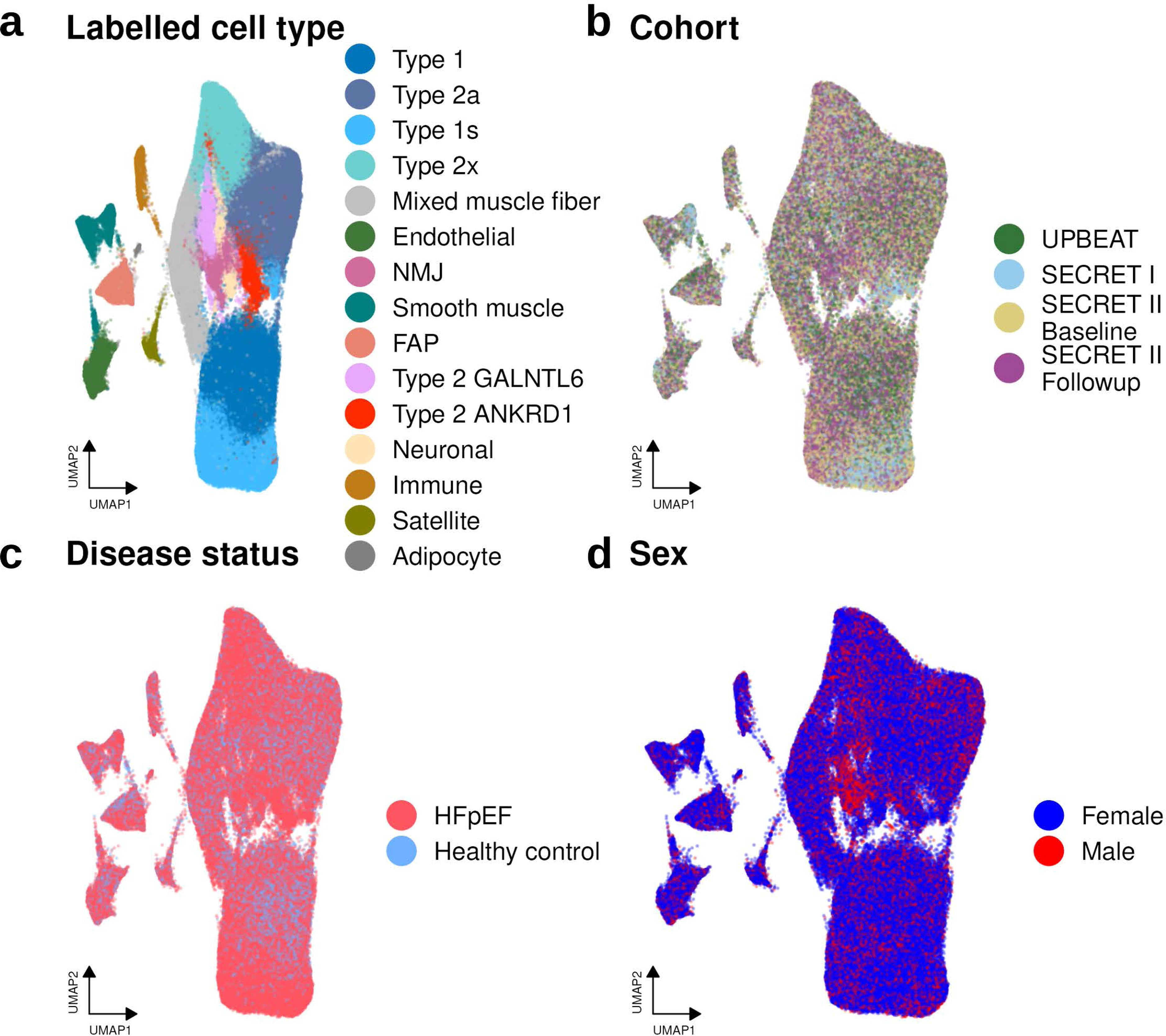
Single-nucleus multiome profiling of human skeletal muscle across HFpEF and healthy control cohorts. UMAP embeddings of nuclei from skeletal muscle biopsies, colored by (top left) annotated cell type, (top right) cohort, (bottom left) disease status, and (bottom right) biological sex.

**Extended Data Fig 2:**
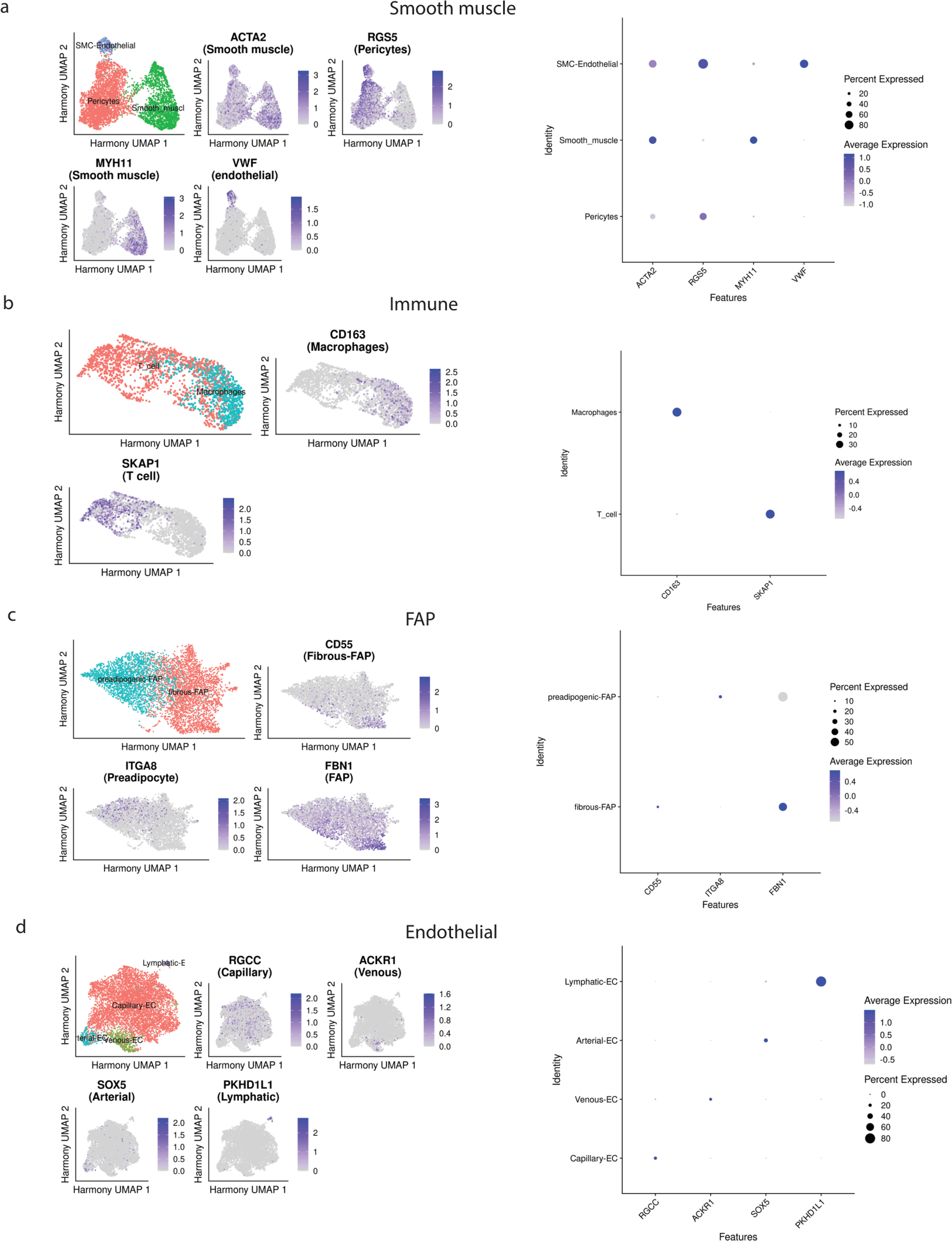
BMI is significantly elevated in HFpEF participants compared to healthy age-matched controls. Box plots show the distribution of BMI across disease groups, with individual data points overlaid. The horizontal line within each box indicates the median, box edges represent the interquartile range (IQR), and whiskers extend to 1.5× the IQR. Significance was assessed by two-sided t-test. *** p < 0.001.

**Extended Data Fig 3:**
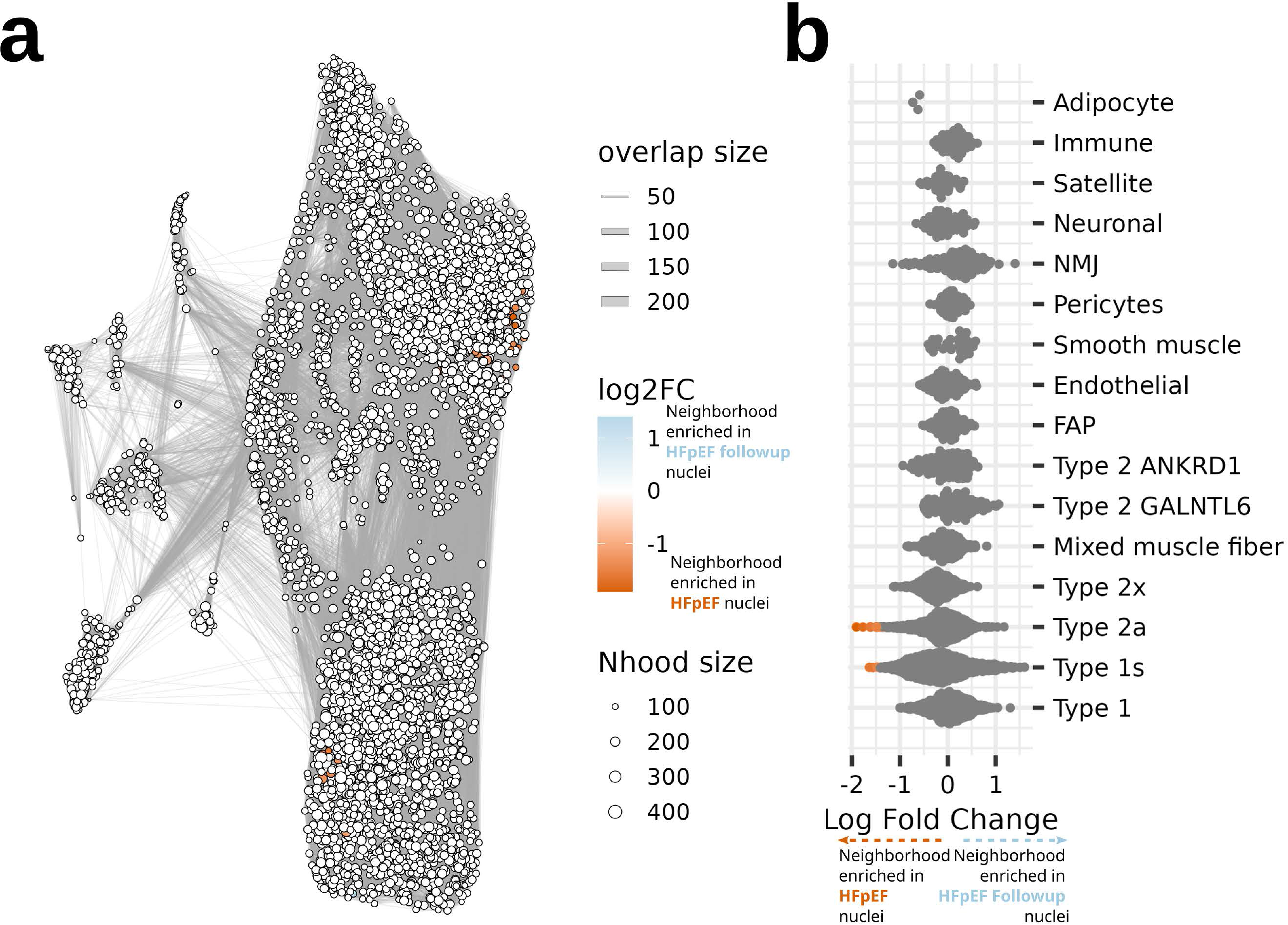
Subclustering identifies key cell subtypes. Subclustering of smooth muscle (a), immune (b), FAP (c), and endothelial (d) showing UMAP plots colored by identified subtypes and key marker genes used to annotated given subtypes (left). Expression specificity of key marker genes depicted by average expression and percent nuclei with expression in the subclusters are shown on the right.

**Extended Data Fig 4:**
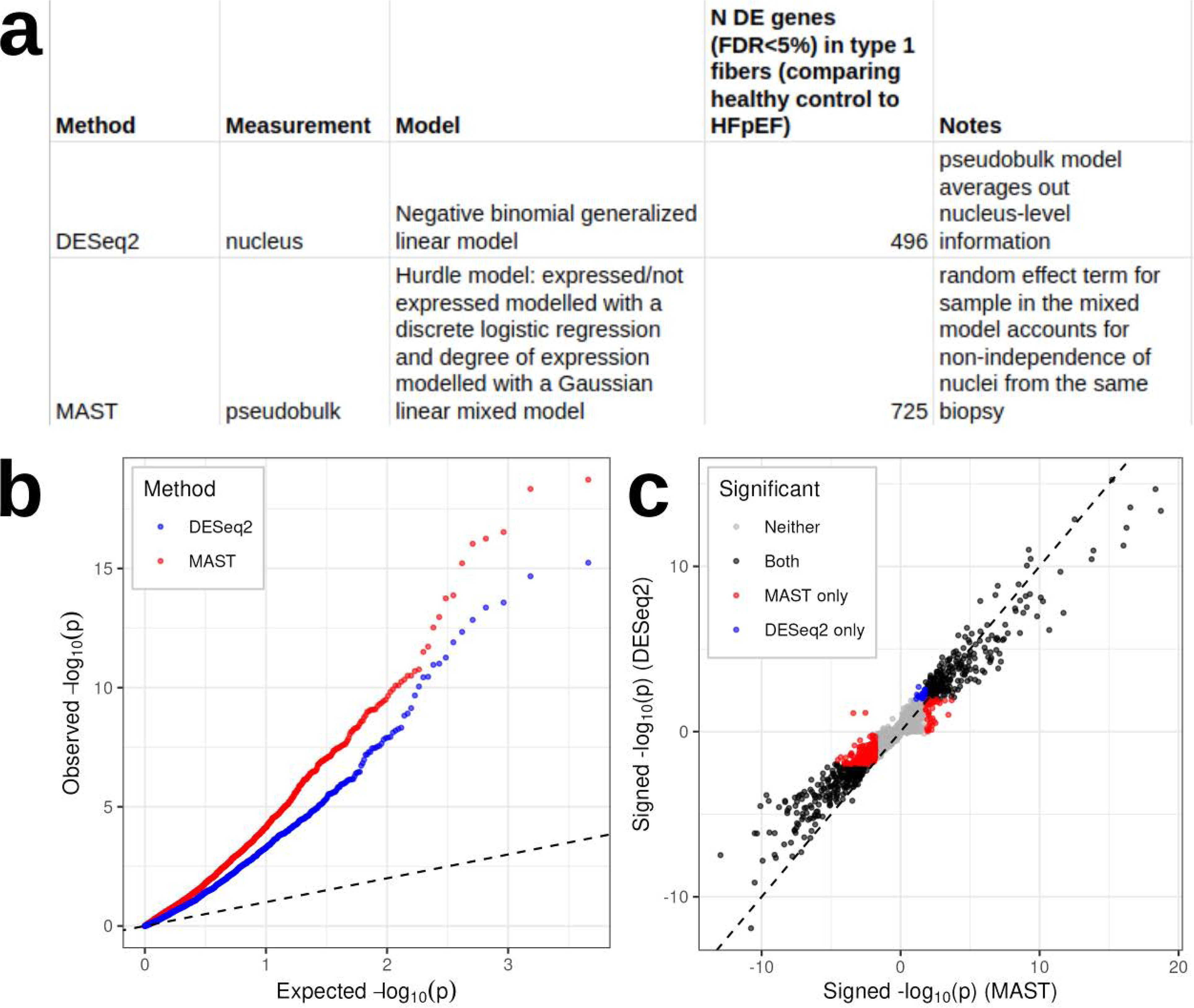
Modest depletion in abundance of Type 1s, Type 2a fibers after supervised diet and exercise intervention. (a) Neighborhood graph of nuclei colored by differential abundance between HFpEF participants and HFpEF participants following a supervised diet and exercise intervention (HFpEF Followup). Orange neighborhoods are significantly (Spatial FDR < 0.05) enriched in HFpEF nuclei; light blue neighborhoods are significantly enriched in HFpEF Followup nuclei. Line thickness between neighborhoods reflects the degree of overlap in nuclei. (f) Beeswarm plot grouping neighborhoods by their majority cell type. Orange neighborhoods are significantly (Spatial FDR < 0.05) enriched in HFpEF nuclei, while light blue neighborhoods are significantly enriched in HFpEF Followup nuclei.

**Extended Data Fig 5:**
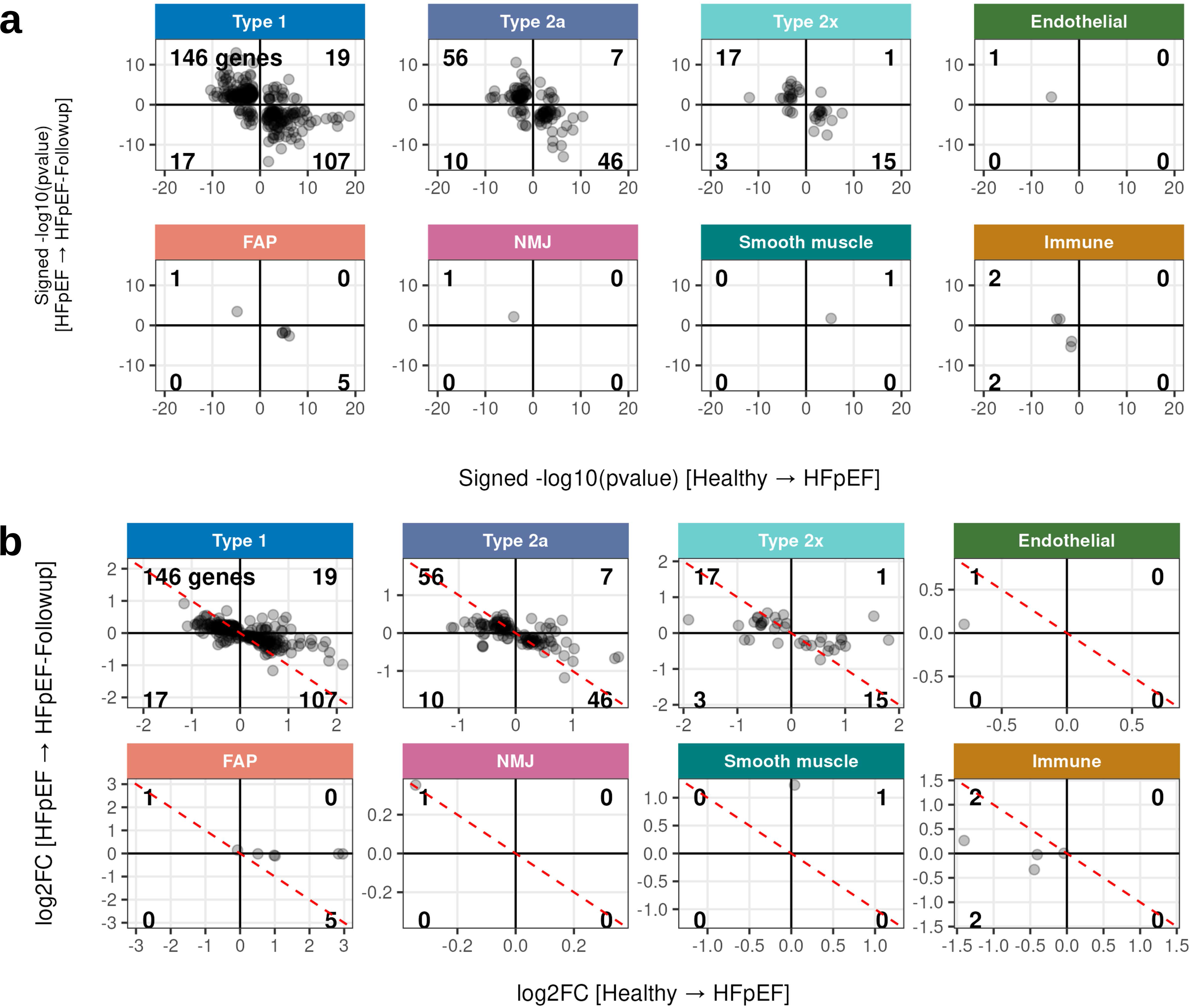
MAST identifies stronger differential expression signals than DESeq2. Each panel shows a scatter plot comparing signed -log10(p-value) estimates from MAST (y-axis) and DESeq2 (x-axis) for all tested genes within the indicated cell type. The sign reflects the direction of effect, with positive values indicating higher expression in healthy controls and negative values indicating higher expression in HFpEF. The red dashed line indicates the identity line (slope = 1).

**Extended Data Fig 6:**
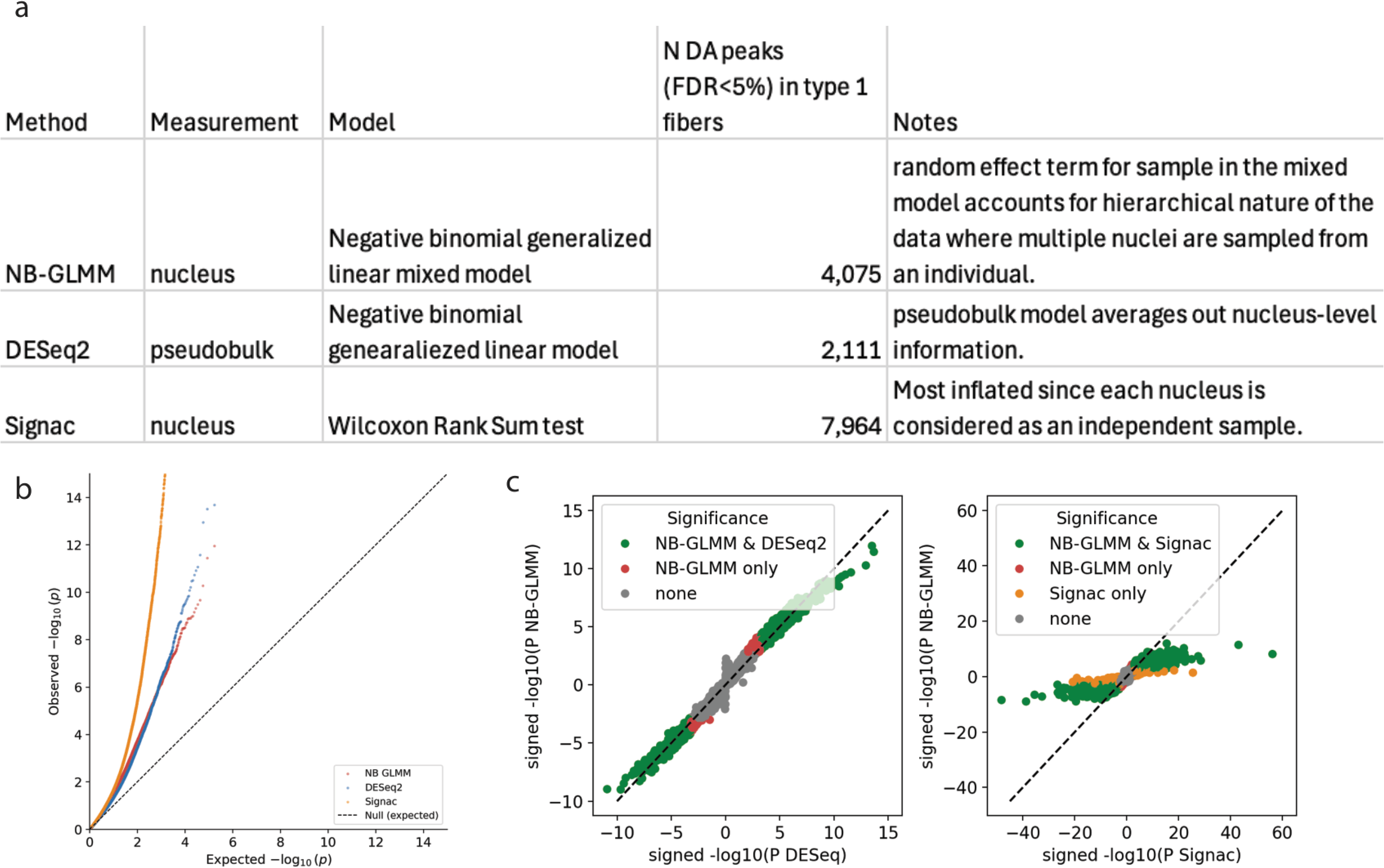
Transcriptional changes induced by HFpEF are partially reversed following treatment in skeletal muscle cell types. (a) Scatter plots comparing signed -log10(p-value) from MAST differential expression testing for two contrasts: Healthy control vs. HFpEF (x-axis) and HFpEF Baseline vs. HFpEF Followup (y-axis) across coarse cell types. The sign reflects the direction of effect, such that genes upregulated in HFpEF relative to healthy controls have positive x-values, and genes upregulated at Followup relative to HFpEF Baseline have positive y-values. (b) Scatter plot comparing MAST log2 fold change (log2FC) estimates for two contrasts: Healthy control vs. HFpEF (x-axis) and HFpEF Baseline vs. HFpEF Followup (y-axis). Positive x-values indicate higher expression in HFpEF relative to healthy controls; positive y-values indicate higher expression at Followup relative to HFpEF Baseline. The red dashed line indicates y = −x. Numbers in each quadrant indicate the count of genes in that quadrant. Only genes meeting a minimum significance threshold in at both contrasts (FDR < 0.05 in one contrast and p < 0.05 in the other) are plotted.

**Extended Data Fig 7:**
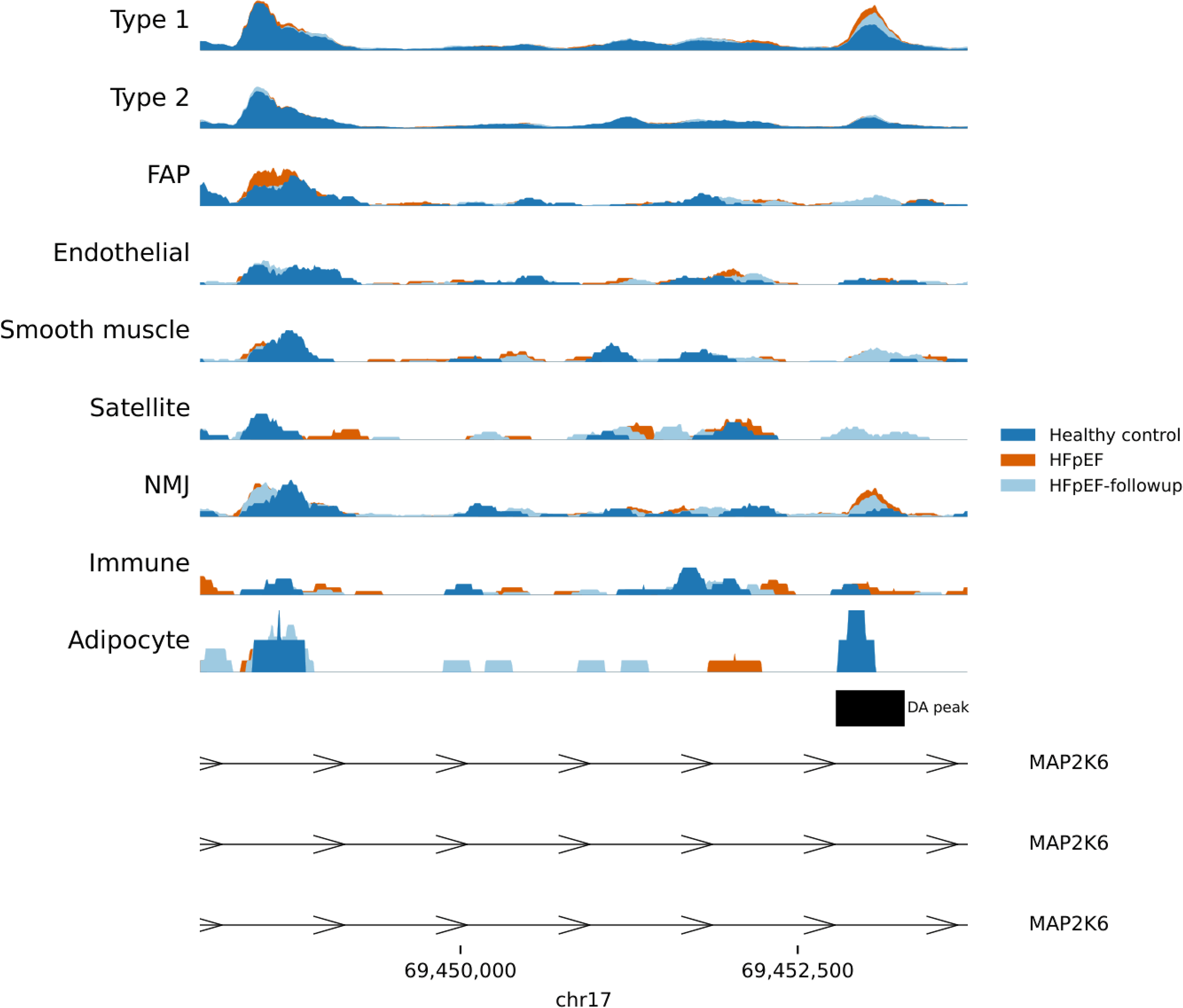
Optimizing DA peak testing. a) Three considered methods for DA peak testing measuring nucleus or pseudobulk level chromatin accessibility. b) q-q plot showing observed p values from the three methods against an expected uniform distribution. c) signed -log10(P) comparisons of DESeq and Signac with NB-GLMM.

**Extended Data Fig 8:**
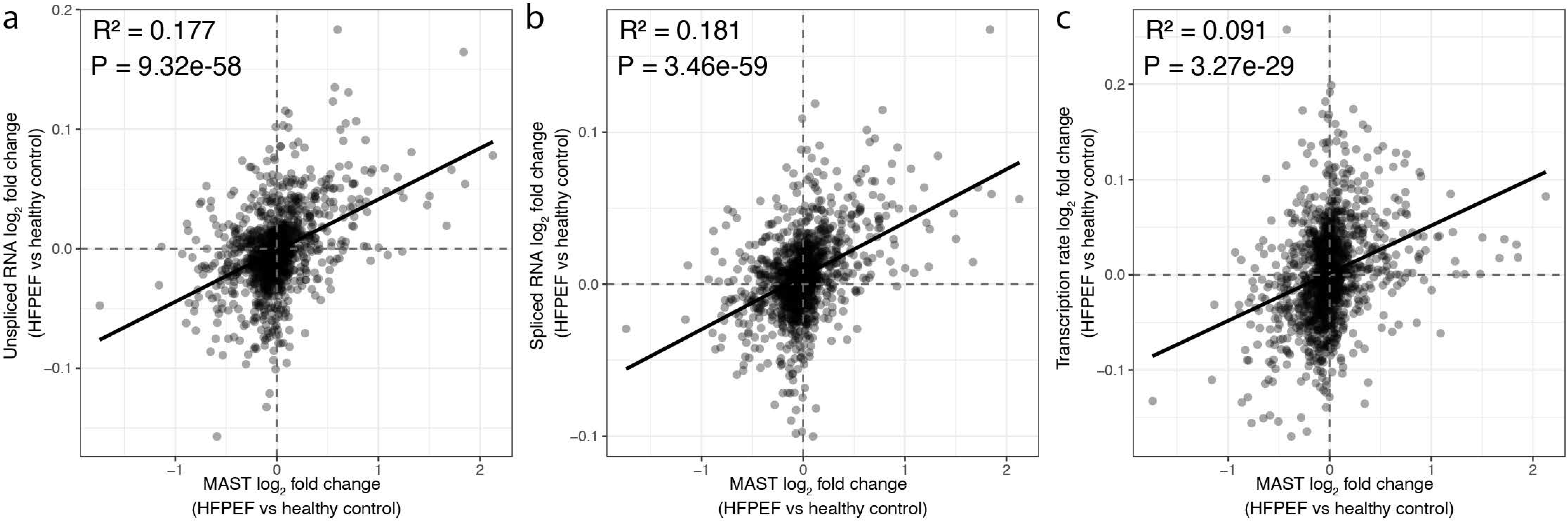
ATACseq signal by disease condition visualization at the *MAP2K6* locus. ATACseq Signal tracks in each cluster by healthy control, HFpEF or HFpEF-followup condition are shown at the *MAP2K6* gene intron on chr17. A DA peak identified in the region is annotated in black.

**Extended Data Fig 9:**
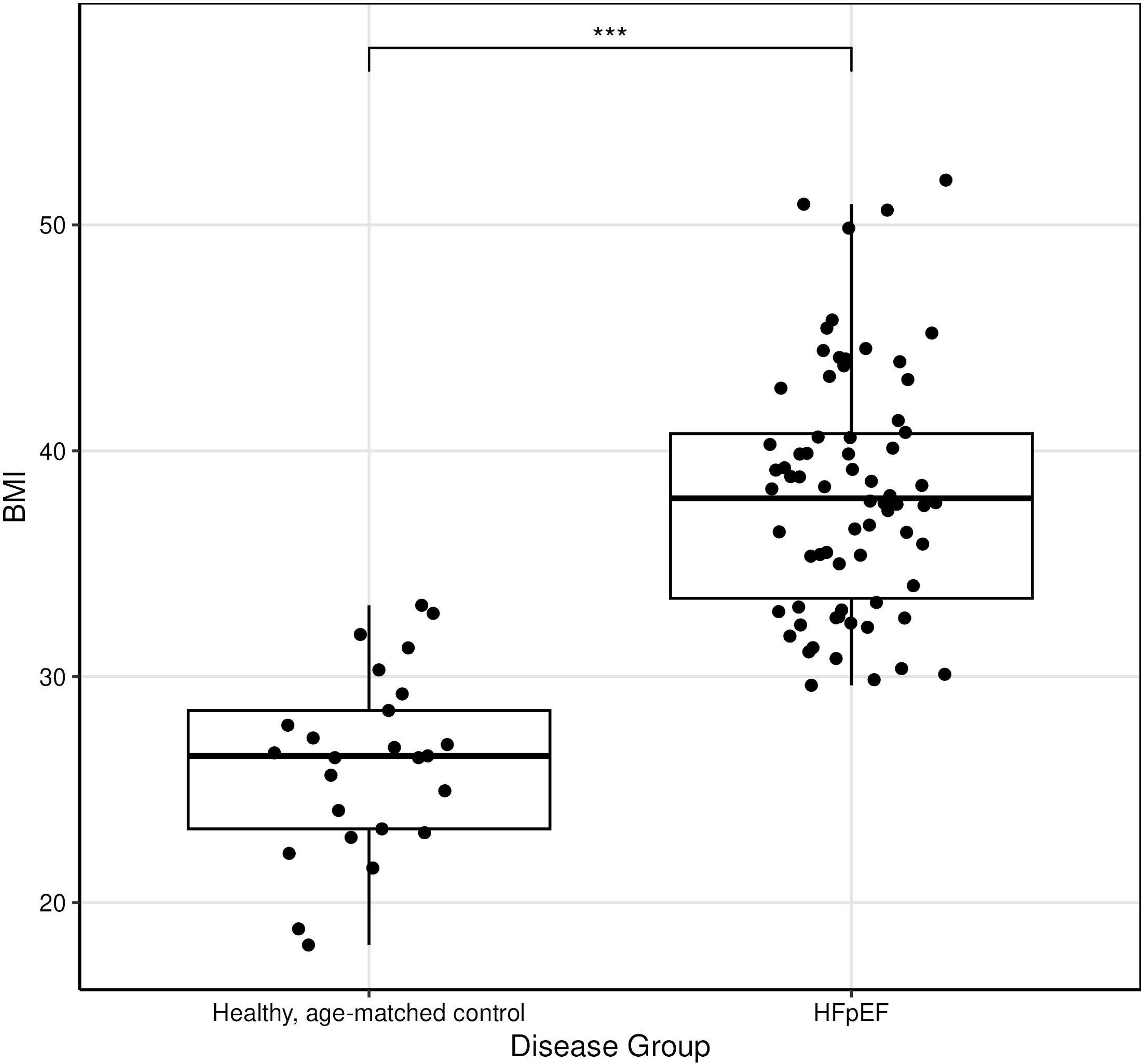
MultiVeloVAE transcription comparison with differential gene expression (MAST). Changes in the MultiVeloVAE estimates for unspliced (a) and spliced (b) RNA abundance and transcription rate (c) were tested for concordance with MAST-derived differential RNA expression data to ensure the two analyses were consistent. Pearson correlation was significant in all cases (shown in top left). The line indicates the best fit.

## List of Supplemental Tables

Supplemental Table S1. Participant metadata characteristics

Supplemental Table S2. DGE results between type 1 nuclei enriched in healthy samples and type 1s nuclei enriched in HFpEF samples

Supplemental Table S3. Gene set enrichment (GO + KEGG) of nominally significantly DE genes

Supplemental Table S4. DGE results between healthy controls and HFpEF samples

Supplemental Table S5. DGE results between HFpEF and HFpEF-followup

Supplemental Table S6. GSE results between healthy controls and HFpEF samples

Supplemental Table S7. GSE results between HFpEF and HFpEF-followup

Supplemental Table S8. GSE results for genes that reverse direction between Healthy → HFpEF and HFpEF → HFpEF-followup

Supplemental Table S9. Motif enrichment in DA peaks

Supplemental Table S10 Nuclei fractions that are coupled or decoupled in Type1s vs Type1 muscle fibers in subjects with HFPEF and healthy controls

Supplemental Table S11 Functional enrichment statistics for genes with differential priming between Type1s and Type1 muscle fibres

Supplemental Table S12 Nuclei fractions that are coupled or decoupled in subjects with HFPEF vs healthy controls in Type1s and Type1 muscle fibers

Supplemental Table S13 Functional enrichment statistics for genes with changes in coupled repression in HFPEF vs healthy controls in Type1s and Type1 muscle fibres

Supplemental Table S14. GWAS enrichment

## List of Supplemental Datasets

Supplemental Dataset 1: Nucleus-QC and cluster annotation

Supplemental Dataset 2: DCA results control vs HFpEF

Supplemental Dataset 3: DCA results HFpEF vs HFpEF-followup

Supplemental Dataset 4: Peaks linked to DE genes (SCENT)

